# Genomic predictive ability for foliar nutritive traits in perennial ryegrass

**DOI:** 10.1101/727958

**Authors:** Sai Krishna Arojju, Mingshu Cao, M. Z. Zulfi Jahufer, Brent A Barrett, Marty J Faville

**Affiliations:** AgResearch Ltd, Grasslands Research Centre, PB 11008, Palmerston North, New Zealand

**Keywords:** genomic selection, genotype-by-environments, heritability, nutritive traits, perennial ryegrass, water soluble carbohydrates

## Abstract

Forage nutritive value impacts animal nutrition, which underpins livestock productivity, reproduction and health. Genetic improvement for nutritive traits has been limited, as they are typically expensive and time-consuming to measure through conventional methods. Genomic selection is appropriate for such complex and expensive traits, enabling cost-effective prediction of breeding values using genome-wide markers. The aims of the present study were to assess the potential of genomic selection for a range of nutritive traits in a multi-population training set, and to quantify contributions of genotypic, environmental and genotype-by-environment (G × E) variance components to trait variation and heritability for nutritive traits. The training set consisted of a total of 517 half-sibling (half-sib) families, from five advanced breeding populations, evaluated in two distinct New Zealand grazing environments. Autumn-harvested samples were analyzed for 18 nutritive traits and maternal parents of the half-sib families were genotyped using genotyping-by-sequencing. Significant (P<0.05) genotypic variation was detected for all nutritive traits and genomic heritability (*h*^*2*^_*g*_) was moderate to high (0.20 to 0.74). G × E interactions were significant and particularly large for water soluble carbohydrate (WSC), crude fat, phosphorus (P) and crude protein. GBLUP, KGD-GBLUP and BayesC genomic prediction models displayed similar predictive ability, estimated by 10-fold cross validation, for all nutritive traits with values ranging from *r* = 0.16 to 0.45 using phenotypes from across two environments. High predictive ability was observed for the mineral traits sulphur (0.44), sodium (0.45) and magnesium (0.45) and the lowest values were observed for P (0.16), digestibility (0.22) and high molecular weight WSC (0.23). Predictive ability estimates for most nutritive traits were retained when marker number was reduced from 1 million to as few as 50,000. The moderate to high predictive abilities observed suggests implementation of genomic selection is feasible for most of the nutritive traits examined. For traits with lower predictive ability, multi-trait genomic prediction approaches that exploit the strong genetic correlations observed amongst some nutritive traits may be useful. This appears to be particularly important for WSC, considered one of the primary constituent of nutritive value for forages.

## 1 Introduction

Perennial ryegrass (*Lolium perenne* L.) from permanent pasture is the major feed component for ruminant production systems in temperate regions of the world. Historically, improvement of annual and seasonal dry matter yield (DMY) have been significant objectives for perennial ryegrass breeding (WILKINS AND HUMPHREYS 2003; WILLIAMS *et al.* 2007; VAN PARIJS *et al.* 2018). Today, seasonal distribution of DMY features as the major component of economic ranking indices developed for this species in New Zealand (Forage Value Index, FVI) (CHAPMAN *et al.* 2017), Australia (LEDDIN *et al.* 2018) and Ireland (Pasture Profit Index, PPI) (MCEVOY *et al.* 2011; MCEVOY *et al.* 2014). Nutritive traits in forages are also important for livestock productivity, maintenance of body weight and for supporting reproduction and health in the grazing animals (WAGHORN AND CLARK 2004). Although there is existing information that demonstrates the importance of nutritive value traits and the potential economic returns from trait improvement, the overall breeding effort for nutritive traits in ryegrass has received considerably less attention than for DMY (SMITH *et al.* 1997). Increased breeding effort for nutritive traits, with validated outcomes for animal productivity, would provide enhanced on-farm value to farmers (JAFARI *et al.* 2003a; CHAPMAN *et al.* 2017).

Compared to other forage grass species, perennial ryegrass is regarded as having relatively high nutritive value, providing a cost effective, nutrient rich feed for ruminant livestock (WILKINS 1991; BAERT AND MUYLLE 2016). Breeding for improved nutritive value in this species has focused principally on higher *in vitro* dry matter (DM) digestibility to enhance energy availability and voluntary intake from grazed pasture (JUNG AND ALLEN 1995). This is a key selection criterion in many ryegrass breeding schemes (CASLER AND VOGEL 1999; EASTON *et al.* 2002; MUYLLE *et al.* 2013), particularly in Europe, where WILKINS AND HUMPHREYS (2003) reported genetic improvement of approximately 10g kg^-1^ per decade for DM digestibility. Breeding to increase water-soluble carbohydrate (WSC) content in ryegrass herbage, one of few reported studies of successful breeding for a nutritive trait in perennial ryegrass (HUMPHREYS 1989a; JONES AND ROBERTS 1991; SMITH *et al.* 1997), has been a major contributor to genetic improvement of digestibility (WILKINS AND HUMPHREYS 2003; MUYLLE *et al.* 2013). More recently, there has been increased emphasis on addressing digestibility through the improvement of fibre degradability *per se*, by targeting changes in the biochemical composition of the cell wall (FAVILLE *et al.* 2010; VAN PARIJS *et al.* 2018).

Minerals and trace elements are essential elements for plant growth and are critical to various biological functions of the plant. In forages, these macro- and micronutrients are also important components of nutritive quality, critical for maintaining livestock health (WAGHORN 2007). For example, metabolic disorders can be caused or contributed to by mineral imbalances in the diet, such as hypomagnesaemia (grass tetany) which is caused by insufficient magnesium and calcium in the diet. Earlier studies have identified genetic variation amongst families (EASTON *et al.* 1997; SMITH *et al.* 1999) or genotypic variation amongst cultivars (CRUSH *et al.* 2018a; CRUSH *et al.* 2018b) for micro- and macronutrients, indicating that breeding for mineral content is a realistic opportunity.

The reduced emphasis on breeding for nutritive traits in forages is affected by a number of factors, including a lack of consensus on specific breeding targets (WHEELER AND CORBETT 1989; CHAPMAN *et al.* 2015), ambiguous evidence for the impact of specific nutritive traits on animal production outcomes (EASTON *et al.* 2002; EDWARDS *et al.* 2007; MCEVOY *et al.* 2011), the confounding influence of environment and genotype × environment (G × E) interactions, and the significant additional resources needed in a breeding program to undertake nutritive trait measurements in large panels of selection candidates (SMITH *et al.* 1997).

Genomic selection (GS), where breeding value for a trait may be cost-effectively predicted for selection candidates using genome-wide markers, was initially proposed for animal breeding by MEUWISSEN *et al.* (2001). In GS, a training population combining phenotypic and genotypic information is used to develop a model that can subsequently be used to predict genomic estimated breeding values (GEBVs) for individuals in a test or selection population that have been genotyped only. In essence, GS replaces the need to phenotype the target trait. GS has been demonstrated in dairy cattle breeding, where the rate of genetic gain was doubled by reducing generation interval from 7 to 2.5 years or from 4 to 2.5 years, depending upon selection strategy (GARCÍA-RUIZ *et al.* 2016). Over the last decade the declining cost of genotyping single nucleotide polymorphisms (SNPs), largely through reduced representation sequencing approaches such as genotyping-by-sequencing (GBS) (ELSHIRE *et al.* 2011), has made this tool feasible for plant breeding. GS is now being applied in major crop species, including wheat (RUTKOSKI *et al.* 2011; POLAND *et al.* 2012; LOPEZ-CRUZ *et al.* 2015; HAYES *et al.* 2017), maize (ZHAO *et al.* 2012; FRISTCHE-NETO *et al.* 2018) and barley (ZHONG *et al.* 2009; LORENZ *et al.* 2012) and is under adoption in forage species, including perennial ryegrass (FÈ *et al.* 2016; GRINBERG *et al.* 2016; BYRNE *et al.* 2017; AROJJU *et al.* 2018; FAVILLE *et al.* 2018; PEMBLETON *et al.* 2018), and alfalfa (ANNICCHIARICO *et al.* 2015; LI *et al.* 2015; BIAZZI *et al.* 2017; JIA *et al.* 2018).

GS can accelerate genetic gain particularly for complex traits, which are controlled by many genes with small effects and for traits which are difficult to measure and expensive (HESLOT *et al.* 2015). GS is therefore a very attractive tool for nutritive traits, given the barriers, described above, to routine integration of nutritive traits into forage breeding programs. The success of GS primarily depends on predictive ability, which is influenced by trait heritability (*h*^*2*^_*n*_), training population size, marker density, extent of linkage disequilibrium (LD) and relatedness between training and test population (DAETWYLER *et al.* 2013). While the heritability of a trait and the extent of LD in a training population cannot be easily optimized, the density of markers and the size and composition of the training population are two factors that can be controlled. Several methods have been developed for genomic prediction and can be broadly classified as whole-genome regression methods (discussed by DE LOS CAMPOS *et al.* (2013)) or machine learning methods (outlined by GONZÁLEZ-CAMACHO *et al.* (2018)). Based on simulation and empirical results, DAETWYLER *et al.* (2013) concluded that genomic best linear unbiased predictor (GBLUP) and Bayesian variable selection methods (BayesB and BayesC) were the benchmark for genomic prediction, as these methods are appropriate for a range of genetic architectures, from traits which are controlled by many genes with small effects (infinitesimal model) to traits with large SNP effects (variable selection model).

The principle aim of the current study was to assess genomic predictive ability for 18 nutritive quality traits, measured in a large multi-population training set in two key New Zealand grazing environments, and to investigate the impact of marker density and of genomic prediction models with different prior assumptions regarding the distribution of SNP effects. The study also provided an opportunity to assess the magnitude of genetic variation and to estimate heritability for a large range of nutritive traits under New Zealand grazing environments.

## 2 Materials and Methods

### 2.1 Plant material and experimental design

The half-sibling (half-sib) families used in this study were derived from five different advanced breeding populations (Pop I − Pop V), which are part of the Grassland Innovation Ltd breeding program. From each population, 102 to 117 plants that tested positive for endophyte infection (*Epichloё festucae* var *lolli*) by immunoblotting (HAHN *et al.* 2003), were polycrossed in isolation during spring 2012 in Palmerston North, New Zealand (FAVILLE *et al.* 2018). Polycrosses were performed separately for each population, without admixing, and seeds from the maternal parents were harvested and cleaned. In total 543 half-sib families were harvested for seed, however only 517 families had sufficient seed (≥ 3.6g) for sowing field trials.

A total of six trials were sown (FAVILLE *et al.* 2018), of which two were used for the current study. These were trials established at Lincoln (Canterbury region, southern New Zealand, 43.38°S 172.62°E; Wakanui silt loam) and Aorangi (Manawatu region, central New Zealand, 40.34°S 175.46°E; Kairanga sandy loam), during the autumn of 2013. The experimental design at each site was row-column with three replicates. Within each replicate, populations were blocked, and families randomized within blocks. Multiple repeated checks (clonal replicates) were also randomly allocated within and across the replicated blocks. Half-sib families were evaluated as a 1m row of plants (referred to from now as plots), by sowing 0.2 g of seed (which is equivalent to 14 kg ha^−1^, if a sward was sown at 7 rows m^−1^). Nitrogen and phosphate fertilizer was applied at the rate of 15-30 kg N ha^−1^ and 8.8 kg P ha^−1^, in late autumn each year (FAVILLE *et al.* 2018).

### 2.2 Phenotypic measurements

Plot harvests were undertaken at Lincoln starting 14 April 2014 and at Aorangi starting 29 April 2014, during the southern hemisphere autumn. At each site a single harvest was undertaken over three days, between 10:30 am and 3:00 pm on each day to minimize the influence of diurnal variation on levels of measured constituents. Split harvesting of populations or replicate blocks over two days was avoided. Plots were cut to a height of approximately 5 cm, above the pseudostem, meaning that only leaf lamina material was harvested. Harvested foliage was placed into micro-perforated plastic bread bags and immediately snap frozen in liquid nitrogen. Samples were subsequently maintained at ca. −80°C on frozen CO_2_ to preserve labile components and then freeze-dried at one of two commercial facilities – Genesis Biolaboratory Ltd (Christchurch, New Zealand) or Horowhenua Freeze-Dry (Levin, New Zealand). Freeze-dried samples were milled to powder through a 1mm sieve and thoroughly mixed to homogenize the sample. Sub-samples were weighed out and transferred to Hill Laboratories (Hamilton, New Zealand) for near-infrared spectroscopy (NIRS) and minerals analysis and to AgResearch (Palmerston North, New Zealand) for analysis of water-soluble carbohydrate (WSC). A total of 3082 samples (n = 1476 from Lincoln and n = 1606 from Manawatu) were provided for analysis. Hill Laboratories provided NIRS data for a range of nutritional traits, as outlined in Table 1. Data for mineral concentrations (Table 1) were based on inductively coupled plasma-optical emission (ICP-OES) analysis of plant material digested with nitric acid: hydrogen peroxide (2:1). Grass tetany ratio was calculated as [K/(Mg + Ca)] using the data provided for the individual minerals. WSC was extracted and quantified as described by HUNT *et al.* (2005). Briefly, 25 mg of milled leaf material was extracted twice with 1mL of 80% ethanol (low-molecular-weight fraction; LMW WSC WSC) and then twice with 1 mL water (high-molecular-weight fraction; HMW WSC WSC), for 30 min at 65°C. Extracts were centrifuged, and supernatants of the respective fractions were analyzed using anthrone as a colorimetric reagent (JERMYN 1956).

**Table 1:**
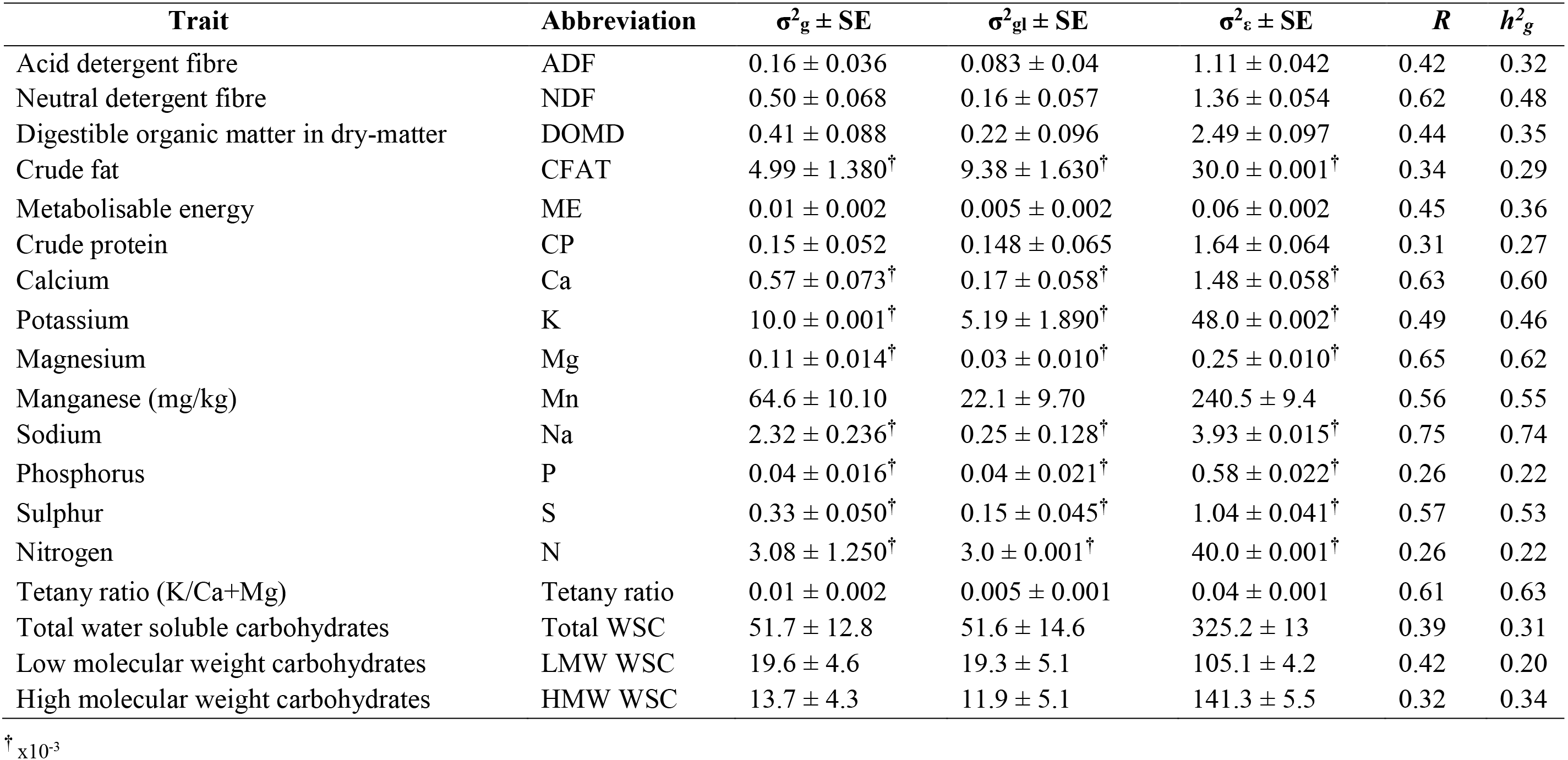
Trait genotypic (σ^2^_g_), genotype-by-location interaction (σ^2^_gl_) and residual error (σ^2^_ε_) variance components and their associated standard errors (SE), repeatability (*R*) and genomic heritability (*h*^*2*^_*g*_), estimated for the range of nutritive traits, among 517 half-sib families of perennial ryegrass evaluated across the two locations in Lincoln and Aorangi. All σ^2^_*g*_ for nutritive traits were significant (*P* < 0.05).

| Trait | Abbreviation | $\sigma^2_g \pm \text{SE}$ | $\sigma^2_{gl} \pm \text{SE}$ | $\sigma^2_\epsilon \pm \text{SE}$ | $R$ | $h^2_g$ |
| --- | --- | --- | --- | --- | --- | --- |
| Acid detergent fibre | ADF | $0.16 \pm 0.036$ | $0.083 \pm 0.04$ | $1.11 \pm 0.042$ | 0.42 | 0.32 |
| Neutral detergent fibre | NDF | $0.50 \pm 0.068$ | $0.16 \pm 0.057$ | $1.36 \pm 0.054$ | 0.62 | 0.48 |
| Digestible organic matter in dry-matter | DOMD | $0.41 \pm 0.088$ | $0.22 \pm 0.096$ | $2.49 \pm 0.097$ | 0.44 | 0.35 |
| Crude fat | CFAT | $4.99 \pm 1.380^\dagger$ | $9.38 \pm 1.630^\dagger$ | $30.0 \pm 0.001^\dagger$ | 0.34 | 0.29 |
| Metabolisable energy | ME | $0.01 \pm 0.002$ | $0.005 \pm 0.002$ | $0.06 \pm 0.002$ | 0.45 | 0.36 |
| Crude protein | CP | $0.15 \pm 0.052$ | $0.148 \pm 0.065$ | $1.64 \pm 0.064$ | 0.31 | 0.27 |
| Calcium | Ca | $0.57 \pm 0.073^\dagger$ | $0.17 \pm 0.058^\dagger$ | $1.48 \pm 0.058^\dagger$ | 0.63 | 0.60 |
| Potassium | K | $10.0 \pm 0.001^\dagger$ | $5.19 \pm 1.890^\dagger$ | $48.0 \pm 0.002^\dagger$ | 0.49 | 0.46 |
| Magnesium | Mg | $0.11 \pm 0.014^\dagger$ | $0.03 \pm 0.010^\dagger$ | $0.25 \pm 0.010^\dagger$ | 0.65 | 0.62 |
| Manganese (mg/kg) | Mn | $64.6 \pm 10.10$ | $22.1 \pm 9.70$ | $240.5 \pm 9.4$ | 0.56 | 0.55 |
| Sodium | Na | $2.32 \pm 0.236^\dagger$ | $0.25 \pm 0.128^\dagger$ | $3.93 \pm 0.015^\dagger$ | 0.75 | 0.74 |
| Phosphorus | P | $0.04 \pm 0.016^\dagger$ | $0.04 \pm 0.021^\dagger$ | $0.58 \pm 0.022^\dagger$ | 0.26 | 0.22 |
| Sulphur | S | $0.33 \pm 0.050^\dagger$ | $0.15 \pm 0.045^\dagger$ | $1.04 \pm 0.041^\dagger$ | 0.57 | 0.53 |
| Nitrogen | N | $3.08 \pm 1.250^\dagger$ | $3.0 \pm 0.001^\dagger$ | $40.0 \pm 0.001^\dagger$ | 0.26 | 0.22 |
| Tetany ratio (K/Ca+Mg) | Tetany ratio | $0.01 \pm 0.002$ | $0.005 \pm 0.001$ | $0.04 \pm 0.001$ | 0.61 | 0.63 |
| Total water soluble carbohydrates | Total WSC | $51.7 \pm 12.8$ | $51.6 \pm 14.6$ | $325.2 \pm 13$ | 0.39 | 0.31 |
| Low molecular weight carbohydrates | LMW WSC | $19.6 \pm 4.6$ | $19.3 \pm 5.1$ | $105.1 \pm 4.2$ | 0.42 | 0.20 |
| High molecular weight carbohydrates | HMW WSC | $13.7 \pm 4.3$ | $11.9 \pm 5.1$ | $141.3 \pm 5.5$ | 0.32 | 0.34 |

$^\dagger \times 10^{-3}$

### 2.3 Statistical models and variance components

Data analyses were performed across the five populations, for individual locations and across the two locations, using the restricted maximum likelihood (REML) method, by fitting a linear mixed model in GenStat (PAYNE *et al.* 2009). Analyses were also performed on the five populations individually, by fitting linear mixed models in DeltaGen (JAHUFER AND LUO 2018). Genotype, G × E interaction, replicates, rows and columns were considered as random effects, whereas location, population and repeated checks were considered as fixed effects. Three different mixed linear models were used: (i) Model 1, to estimate genotypic variance components, pooling all five populations, all 517 families together, within individual locations; (ii) Model 2, for estimating genotypic variance components and interactions of family and location, pooling all five populations, across locations; and (iii) Model 3, for estimating genetic variance and G × E interactions, among half-sib families within individual populations, across locations.

Model 1: Mixed model for individual locations.

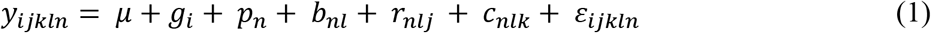

*y*_*ijkln*_ is the phenotypic value measured on half-sib family *i* in row *j* and column *k* of replicate *l* nested within population *n*, and *i* = 1, …, *n*_*g*_, *j* = 1, …, *n*_*r*_, *k* = 1, …, *n*_*c*_, *l* = 1, …, *n*_*b*_, *m* = 1, …, *n*_*s*_, *n* = 1, …, *n*_*p*_, where *g*, *r*, *c*, *b*, and *p* are half-sib families, rows, columns, replicates and populations respectively. Where, *μ* is the overall mean; *g*_*i*_ is the random effect of half-sib family *i*, 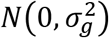; *p*_*n*_ is the fixed effect of population *n*; *b*_*nl*_ is the random effect of replicate *l* in population *n*, 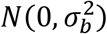; *r*_*nlj*_ is the random effect of row *j* within replicate *l* of population *n*, 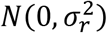; *c*_*nlk*_ is the random effect of column *k* within replicate *l* of population *n*, 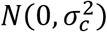; *ε*_*ijkln*_ is the residual effect of half-sib family *i* in row *r* and column *c* of replicate *b* of population *n*, 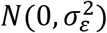.

Model 2: Mixed model for across locations.

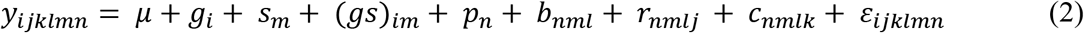

*y*_*ijklmn*_ is the phenotypic value measured on half-sib family *i* in row *j* and column *k* of replicate *l* nested in location *m* within population *n*, and *i* = 1, …, *n*_*g*_, *j* = 1, …, *n*_*r*_, *k* = 1, …, *n*_*c*_, *l* = 1, …, *n*_*b*_, *m* = 1, …, *n*_*s*_, *n* = 1, …, *n*_*p*_, where *g*, *r*, *c*, *b*, *s* and *p* are half-sib families, rows, columns, replicates, locations and populations respectively. In the equation, *μ* is the overall mean; *g*_*i*_ is the random effect of half-sib family *i*, 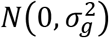; *s*_*m*_ is the fixed effect of location *m*; (*gs*)_*im*_ is the random effect of interaction between half-sib family *i* and location *m*, 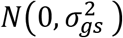; *p*_*n*_ is the fixed effect of population *n*; *b*_*nml*_ is the random effect of replicate *l* within location *m* in population *n*, 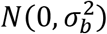; *r*_*nmlj*_ is the random effect of row *j* within replicate *l* in location *m* of population *n*, 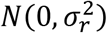; *c*_*nmlk*_ is the random effect of column *k* within replicate *l* in location *m* of population *n*, 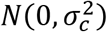; *ε*_*ijklmn*_ is the residual effect of half-sib family *i* in row *r* and column *c* of replicate *b* in location *m* of population *n*, 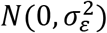.

Model 3: Mixed model for individual populations.

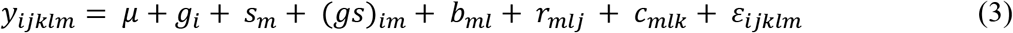

*y*_*ijklm*_ is the phenotypic value measured on half-sib family *i* in row *j* and column *k* of replicate *l* nested in location *m*. In the equation, *μ* is the overall mean; *g*_*i*_ is the random effect of half-sib family *i*, 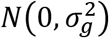; *s*_*m*_ is the fixed effect of location *m*; (*gs*)_*im*_ is the random effect of interaction between half-sib family *i* and location *m*, 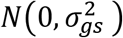; *p*_*n*_ is the fixed effect of population *n*; *b*_*ml*_ is the random effect of replicate *l* within location *m*, 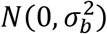; *r*_*mlj*_ is the random effect of row *j* within replicate *l* in location *m*, 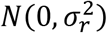; *c*_*mlk*_ is the random effect of column *k* within replicate *l* in location *m*, 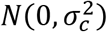; *ε*_*ijklmn*_ is the residual effect of half-sib family *i* in row *r* and column *c* of replicate *b* in location *m*, 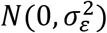.

The variance components estimated based on the mixed model analysis were used to calculate repeatability (Model 2) (FALCONER 1960) and narrow sense heritability (Model 3) for each trait. Repeatability was based on genotypic variance estimated across five populations, whereas narrow-sense heritability is based on additive genetic variance among half-sib families within each population. Repeatability and narrow sense heritability, on a family mean basis, were estimated using the equation:

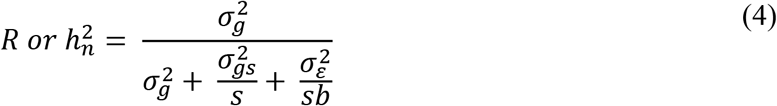

Where, *R* and 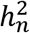 are repeatability and narrow-sense heritability. For repeatability, 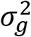 was the genotypic variance among all the 517 half-sib families. In the estimation of narrow-sense heritability, 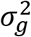 was the estimated additive genetic variation among half-sib families within a specific population, 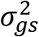 is the variance associated with G × E interaction and 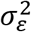 is the variance of residuals.

### 2.4 Genotypic and phenotypic correlation

The genotypic correlation among traits was estimated as proposed by FALCONER (1960). Multivariate analysis of variance (MANOVA) was performed in DeltaGen (JAHUFER AND LUO 2018), using the multivariate analysis option, to estimate variance and covariance among traits:

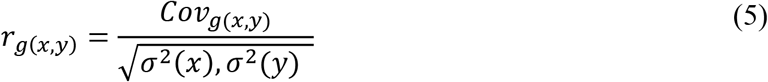

Where, *Cov*_*g*(*x*,*y*)_ is the genotypic covariance between trait *x* and *y*; *σ*^2^(*x*) is the variance associated with trait *x*, and *σ*^2^(*y*) is the variance associated with trait *y*. Phenotypic correlation was performed in DeltaGen (JAHUFER AND LUO 2018) using the best linear unbiased predictors (BLUPS) estimated based on Model 2.

### 2.5 Genotyping and genomic heritability

All maternal parents of the 517 half-sib families were genotyped using a GBS approach described in FAVILLE *et al.* (2018), following the protocol proposed by ELSHIRE *et al.* (2011). Briefly, a reference ryegrass genome assembly was constructed using scaffolds from a published ryegrass assembly (BYRNE *et al.* 2015). Scaffolds were aligned to the barley genome using Lastz version 7.0.1 (HARRIS 2007) from Geneious 8 (https://www.geneious.com/, (KEARSE *et al.* 2012)) with default parameters. Demultiplexing of sequencing reads was performed using the TASSEL 5.0 GBS pipeline (GLAUBITZ *et al.* 2014) and initial quality control was based on read count statistics. The quality GBS tags were aligned to the reference genome using Bowtie2 (LANGDON 2015). Genotype calling was performed using TASSEL GBS pipeline to obtain 1,093,464 SNPs, after filtering for maximum missing SNPs per site (50%), minor allele frequency (> 0.05) and read depth (> 1) using VCF tools (DANECEK *et al.* 2011). Genotyped 1,093,464 SNPs were exported and filtered for Hardy-Weinberg disequilibrium (HWdiseq > −0.05). The resulting 1,023,011 SNPs, with a mean read depth of 2.98, were used to compute a genomic relationship matrix (KGD matrix) based on protocol proposed by DODDS *et al.* (2015). The KGD matrix was used for genomic predictive modelling. Population structure was previously analyzed using multi-dimensional scaling based on genomic relationship matrix (see Figure 1 in FAVILLE *et al.* (2018))

**Figure 1:**
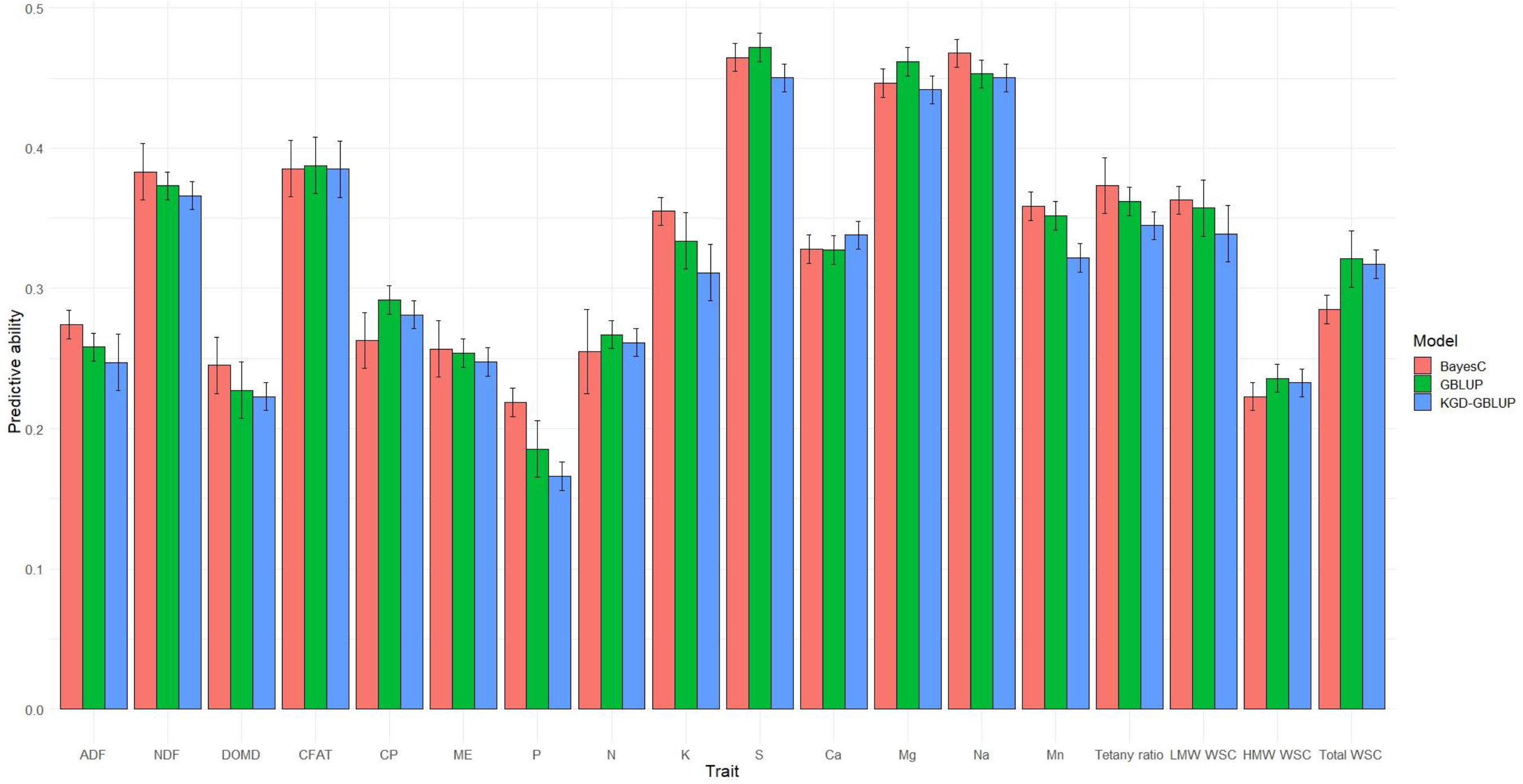
Predictive ability (Pearson correlation coefficient between observed and predicted values) for nutritive traits and their associated standard deviation, assessed using three genomic prediction models (BayesC, KGD-GBLUP and GBLUP), based on adjusted means (BLUP’s) measured among five populations across two locations.

Genomic heritability (*h*^*2*^_*g*_) was calculated using Eq. 4, based on variance components estimated using the mixed model proposed in Eq. 2. In the model, the KGD matrix was fitted as variance-covariance among genotypes (DE LOS CAMPOS *et al.* 2015) and the genetic variance was calculated as proportion of variance explained by regressing markers on phenotypes. The model was fitted in ASreml-R (BUTLER *et al.* 2009).

### 2.6 Genomic prediction modelling

Three whole-genome regression methods, with two different prior assumptions regarding the distribution of marker effects, were used for generating GEBVs. The first method was a univariate linear mixed model, called GBLUP (GODDARD *et al.* 2011) in which markers effects were assumed to have equal variance. The linear model can be expressed follows:

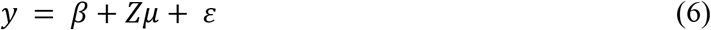

Where *y* is the vector of BLUP values of the trait, *β* is the vector of grand mean, *Z* is the design matrix associated with random marker effects *μ*, with 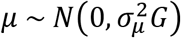, in which *G* is the additive genetic relationship matrix, and 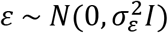, in which *I* is the identity matrix. The *G* matrix was calculated based on the method proposed by VANRADEN (2008); ENDELMAN AND JANNINK (2012) using A.mat function in rrBLUP package (ENDELMAN 2011).

The second method is a variant of GBLUP method with KGD matrix as *G* in the linear mixed model. The GBLUP and KGD-GBLUP models were fitted using the rrBLUP package (ENDELMAN 2011), implemented through R programming language (R CORE TEAM 2017).

The third method was BayesC (HABIER *et al.* 2011), in which markers effects can depart from normality, that is, large variances are allowed for markers with larger effects.

The model is expressed as follows:

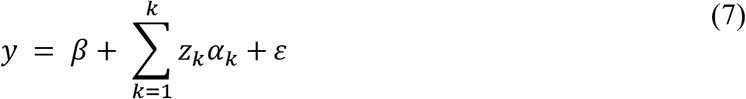

Where *y* is the vector of BLUP values of the trait, *β* is the vector of grand mean, *k* is the number of makers, *Z*_*k*_ is the vector of genotypes at marker *k*, *α*_*k*_ is the additive effect of the marker, and *ε* is the vector of residual effects with a normal distribution 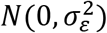. The BayesC model was implemented through R programming using the BGLR package (PÉREZ AND DE LOS CAMPOS 2014), with the number of burn-ins set to 2,000, total number of iterations set to 10,000, and other parameters set to default (PÉREZ AND DE LOS CAMPOS 2014).

The predictive ability of the models based on data from the composite training population was assessed by a ten-fold cross validation approach. For each cross validation, randomized data were divided into ten equal parts, of which nine parts (training set) were used to train the model and to predict GEBVs in the remaining one part of the data (test set). Randomization of the complete data set was repeated five times and the mean of the five iterations was reported as the predictive ability of the model (FAVILLE *et al.* 2018).

### 2.7 Evaluating predictions in individual populations

As the overall training population is a composite of 517 individuals and their corresponding half-sib families from five discrete breeding populations, the predictive ability of the prediction models was also assessed within each individual population using KGD-GBLUP. A random 50% of individuals was selected from within each population (Pop I – Pop V; total = 255 individuals) as a training set in order to represent each population equally. Using this set of 255 individuals to train the model, GEBVs were then predicted in the remaining 50% of Pop I and the mean correlation of 500 iterations was considered as the predictive ability for this population. This approach was likewise extended to each of the other four populations.

### 2.8 Optimising marker density

To evaluate the minimum number of markers needed to achieve maximum predictive ability for each nutritive trait, a random set of markers ranging from 1,093,464 (100%, unfiltered) to 1,093 (0.1%) in 10 steps were obtained from the training population. Using each set of randomly selected markers, a *G* matrix was computed based on the method proposed by VANRADEN (2008) using the rrBLUP package (ENDELMAN 2011). Considering the computational load, KGD method was not extended to randomly selected markers, to construct *G* matrix. FAVILLE *et al.* (2018) reported broadly similar predictive ability for DMY in this training population, when *G* matrices based on DODDS *et al.* (2015) and VANRADEN (2008) were compared. The *G* matrix was used in a GBLUP model to estimate predictive ability for each randomly chosen marker set. The predictive ability was assessed via Monte-Carlo cross validations with 500 iterations, where 80% of the data were used to train the model (training set) and 20% to predict the GEBVs (test set).

## 3 Results

### 3.1 Variance components, repeatability, and genomic heritability

There was significant (P<0.05) genotypic variation among 517 half-sib families from five populations for all traits, based on mean performance across the two locations, Lincoln and Aorangi (Table 1, Supplementary Table S1 and S2). There were also significant (P<0.05) G × E interactions for all the traits, indicating a relative change in ranking among the 517 half-sib families between the two locations. There was a high genotypic correlation (*r* = 0.93) between *R* and *h*^*2*^_*g*_ in the across-location dataset and these ranged from a low of 0.26 (*R*) and 0.22 (*h*^*2*^_*g*_) for traits N and P to a high of 0.75 (*R*) and 0.74 (*h*^*2*^_*g*_) for Na (Table 1) across the two locations. Genotypic correlation between *R* and *h*^*2*^_*g*_ was slightly lower in Aorangi (*r* = 0.85) compared with Lincoln (*r* = 0.93). Because of the high correlation between *R* and *h*^*2*^_*g*_ and because *h*^*2*^_*g*_ captures marker-based additive variance, from here on results for *h*^*2*^_*g*_ only are reported and discussed. Overall, *h*^*2*^_*g*_ estimated within a location was substantially higher at the Aorangi site than Lincoln (mean of all traits *h*^*2*^_*g*_ = 0.62 and 0.43, respectively) (Supplementary Table S1 and S2), with values from the across-location analysis (*h*^*2*^_*g*_ = 0.42) lying between those of Lincoln and Aorangi. Traits with low *h*^*2*^_*g*_ tended to have relatively large G × E, whereas those with high *h*^*2*^_*g*_ had a low G × E interaction components (Table 1). Variance component analysis within the two locations (Lincoln and Aorangi) indicated significant (P<0.05) genotypic variation for all 18 nutritive traits. Differences in additive variance were observed for the same trait amongst the five populations in the across location dataset (Supplementary Table S3-S7). For example, additive genetic variance was non-significant (P > 0.05) for ADF, NDF and DOMD in Pop I & II, but was significant for these traits in Pop III – V (Supplementary Table S3-S7). Similar observations can be made for all of the analyzed traits, with no population showing significant (P < 0.05) additive genetic variance component for all 18 traits. Amongst the five populations, Pop I had significant additive variance for only 42% of traits (8 traits out of 19) while for Pop V that number was 84%, with the remaining populations intermediate to these at 58 – 68% (Supplementary Table S3-S7).

### 3.2 Correlation among traits

Genotypic and phenotypic correlation coefficients for all nutritive quality traits are shown in Tables 2 and S8, respectively. Strong, positive genotypic correlation was observed between fibre measures ADF and NDF and these in turn were negatively correlated with energy traits including ME, DOMD and WSC (Tables 2 and Supplementary Table S8). A positive genotypic correlation was estimated for both LMW WSC and total WSC with DOMD, however, a weak positive correlation was found between HMW WSC and DOMD. A strong negative genotypic correlation was observed for both ADF and NDF with both LMW WSC and total WSC. A moderate genotypic correlation was observed between fibre traits (ADF and NDF) and minerals traits including K, Mg and Mn (positive), P and Ca (negative).

**Table 2:** Genotypic correlations amongst a range of nutritive quality traits measured from 517 half-sib families, estimated using data from across two locations (Lincoln and Aorangi).

|  | ADF | NDF | DOMD | CFAT | CP | ME | P | N | K | S | Ca | Mg | Na | Mn | Tetany ratio | LMW WSC | HMW WSC | Total WSC |
| --- | --- | --- | --- | --- | --- | --- | --- | --- | --- | --- | --- | --- | --- | --- | --- | --- | --- | --- |
| ADF | 1 | 0.84 | -0.88 | 0.12 | -0.65 | -0.87 | -0.51 | -0.69 | 0.47 | -0.26 | -0.72 | 0.20 | 0.03 | 0.30 | 0.61 | -0.49 | -0.04 | -0.35 |
| NDF |  | 1 | -0.77 | -0.05 | -0.53 | -0.77 | -0.43 | -0.54 | 0.60 | -0.09 | -0.79 | 0.41 | -0.23 | 0.54 | 0.67 | -0.60 | -0.38 | -0.59 |
| DOMD |  |  | 1 | 0.05 | 0.36 | 1.00 | 0.57 | 0.43 | -0.29 | 0.15 | 0.46 | -0.45 | -0.01 | -0.55 | -0.32 | 0.73 | 0.21 | 0.60 |
| CFAT |  |  |  | 1 | -0.10 | 0.03 | 0.73 | -0.06 | 0.02 | -0.61 | 0.00 | -0.67 | 0.32 | -0.57 | 0.14 | 0.41 | 0.59 | 0.55 |
| CP |  |  |  |  | 1 | 0.36 | 0.27 | 0.99 | -0.49 | 0.06 | 0.68 | 0.14 | 0.02 | 0.10 | -0.65 | -0.19 | 0.11 | -0.08 |
| ME |  |  |  |  |  | 1 | 0.55 | 0.44 | -0.29 | 0.18 | 0.46 | -0.42 | 0.01 | -0.55 | -0.32 | 0.72 | 0.22 | 0.59 |
| P |  |  |  |  |  |  | 1 | 0.34 | -0.15 | -0.20 | 0.29 | -0.62 | 0.13 | -0.53 | -0.13 | 0.58 | 0.33 | 0.55 |
| N |  |  |  |  |  |  |  | 1 | -0.45 | 0.08 | 0.65 | 0.10 | 0.01 | 0.07 | -0.60 | -0.13 | 0.11 | -0.04 |
| K |  |  |  |  |  |  |  |  | 1 | 0.25 | -0.69 | 0.34 | -0.61 | 0.45 | 0.92 | -0.29 | -0.55 | -0.46 |
| S |  |  |  |  |  |  |  |  |  | 1 | -0.08 | 0.55 | -0.50 | 0.47 | 0.10 | -0.19 | -0.67 | -0.44 |
| Ca |  |  |  |  |  |  |  |  |  |  | 1 | -0.07 | 0.35 | -0.34 | -0.89 | 0.31 | 0.34 | 0.37 |
| Mg |  |  |  |  |  |  |  |  |  |  |  | 1 | -0.43 | 0.82 | 0.09 | -0.75 | -0.66 | -0.82 |
| Na |  |  |  |  |  |  |  |  |  |  |  |  | 1 | -0.58 | -0.47 | 0.32 | 0.73 | 0.55 |
| Mn |  |  |  |  |  |  |  |  |  |  |  |  |  | 1 | 0.29 | -0.89 | -0.83 | -1.00 |
| Tetany ratio |  |  |  |  |  |  |  |  |  |  |  |  |  |  | 1 | -0.20 | -0.37 | -0.31 |
| LMW WSC |  |  |  |  |  |  |  |  |  |  |  |  |  |  |  | 1 | 0.50 | 0.92 |
| HMW WSC |  |  |  |  |  |  |  |  |  |  |  |  |  |  |  |  | 1 | 0.80 |
| Total WSC |  |  |  |  |  |  |  |  |  |  |  |  |  |  |  |  |  | 1 |

### 3.3 Predictive ability for nutritive traits

Predictive ability for all nutritive traits was evaluated using GBLUP, KGD-GBLUP and BayesC genomic prediction models, and the results are summarized in Figure 1 as the Pearson correlation coefficient between observed (adjusted means) and predicted values. There were no significant differences (P > 0.05) in terms of predictive ability between GBLUP, KGD-GBLUP and BayesC across all nutritive traits (Figure 1). Although slight differences can be noted from the Figure 1, no single statistical approach consistently gave higher predictive ability for all nutritive traits. Because the results from the three models were largely indistinguishable, from here on results from KGD-GBLUP are only reported and discussed. Using the adjusted phenotypic trait means (BLUPs) estimated across both locations, predictive ability for all traits was positive and was strongly correlated with *h*^*2*^_*g*_ (*r* = 0.65). The highest predictive ability observed was for Na and S (both *r* = 0.45), followed by CFAT (0.38) (Figure 1). The lowest predictive ability was noted for P (0.16), followed by DOMD with a value of 0.22 (Figure 1). The bias (slope of regression) of the model for all nutritive traits was around 1, meaning unbiased estimates were obtained by regressing GEBVs on adjusted means (BLUPs) (Supplementary Table S9).

Predictive ability of models based on phenotypic means from Lincoln only (location-specific predictive ability) was negative to low and showed a very high correlation with *h*^*2*^_*g*_ (*r* = 0.93) (Supplementary Table S1). The highest predictive ability was obtained for Na (0.35), similar to the across locations analysis, and the lowest predictive ability was for ADF with a negative accuracy of −0.06. Predictive ability of models using phenotypic data from Aorangi were generally higher than both the Lincoln and across-location models (Supplementary Table S2) and the correlation between *h*^*2*^_*g*_ and predictive ability was 0.67. In this dataset the highest predictive ability was for HMW WSC (0.56) and lowest predictive ability was for Ca (0.16) (Supplementary Table S2).

In terms of different trait categories, for the measures of fibre content, ADF and NDF, predictive ability of the across-location models was moderate, at 0.24 and 0.36 respectively. There was a strong effect of location on these traits, with moderate predictive ability at Aorangi (ADF = 0.29 and NDF = 0.35) whereas at Lincoln, the predictive ability was almost zero for NDF (0.02) and negative for ADF (−0.06) (Supplementary Table S1 and S2).

The traits DOMD, CFAT, WSC (LMW, HMW and total) and ME were grouped as energy traits in this study. Predictive ability for energy traits in the across location analysis was generally low to moderate, with CFAT (0.38) and LMW WSC (0.34) the highest, and DOMD (0.22) and HMW WSC (0.23) low (Figure 1). As with the fibre traits, the ranking of predictive ability for CFAT varied by environment and was highest in Lincoln and in across-location analysis, whereas predictive ability for CFAT ranked fourth highest in Aorangi. By contrast, the predictive ability estimated for DOMD was ranked similarly (fifth highest) for Lincoln and Aorangi.

The predictive ability of genomic prediction models for mineral traits assessed in this study was generally high, with Mg, Na and S consistently ranked highest in terms of predictive ability within the two locations (Lincoln and Aorangi) and in across-location analysis. The lowest *h*^*2*^_*g*_ was observed for P, which was reflected in the predictive ability of prediction models for Lincoln and across-location analysis. Models for tetany ratio ([K/(Ca+Mg)]), a predictor of hypomagnesaemia risk in livestock, had a predictive ability of 0.34 across locations, 0.29 at Lincoln and 0.18 at Aorangi.

The measures CP and N are both indicative of protein content, with crude protein a derivate of measured N, obtained by multiplying N by a conversion factor of 6.25 (WAGHORN 2007), hence predictive ability estimated within and across locations was highly similar for both the traits. Predictive ability for these traits was low to moderate, at 0.28 (CP) and 0.26 (N) in the across location analysis, 0.14 for both traits at Lincoln and 0.20 and 0.21 for CP and N at Aorangi.

Genotyping efficiency impacts the design and overall cost of implementing GS in a breeding program. To investigate the minimum number of SNP markers needed to achieve maximum predictive ability within the current dataset, random marker sets with varying numbers of SNPs were used to build genomic prediction models for all nutritive traits, using the across locations dataset. For all nutritive traits, a steady decline in predictive ability was observed from 100% (1,093,464) to 0.5% (5,467) markers and a rapid decrease in predictive ability was noted from 0.5% to 0.1% (1,093) (Figure 2 and Supplementary Table S9). Overall, reducing the marker number to 5% (54,673) of the total available SNPs had minimal impact on overall predictive ability (Figure 2 and Supplementary Table S9). Further reductions in marker number resulted in losses in predictive ability, the extent of which varied by trait (Supplementary Table S9). For example, with 10,934 markers (1% of the total dataset) the predictive ability for LMW WSC, HMW WSC and total WSC decreased by 3%, 7% and 4%, respectively compared to the total dataset (100%) (Figure 2). At 1,093 markers (0.1%) the predictive ability for these traits declined further although the absolute values were still positive, at 0.31 for LMW WSC, 0.18 for HMW WSC and 0.26 for total WSC (Figure 2). The decay in predictive ability was typically highest for those traits which had low *h*^*2*^_*g*_ and low predictive ability under the full SNP dataset. For example, between the highest and lowest marker number datasets there was a 36% decrease in predictive ability for P (*h*^*2*^_*g*_ = 0.22), while for S (*h*^*2*^_*g*_ = 0.53) there was a 14% decrease in predictive ability (Supplementary Table S9).

**Figure 2:**
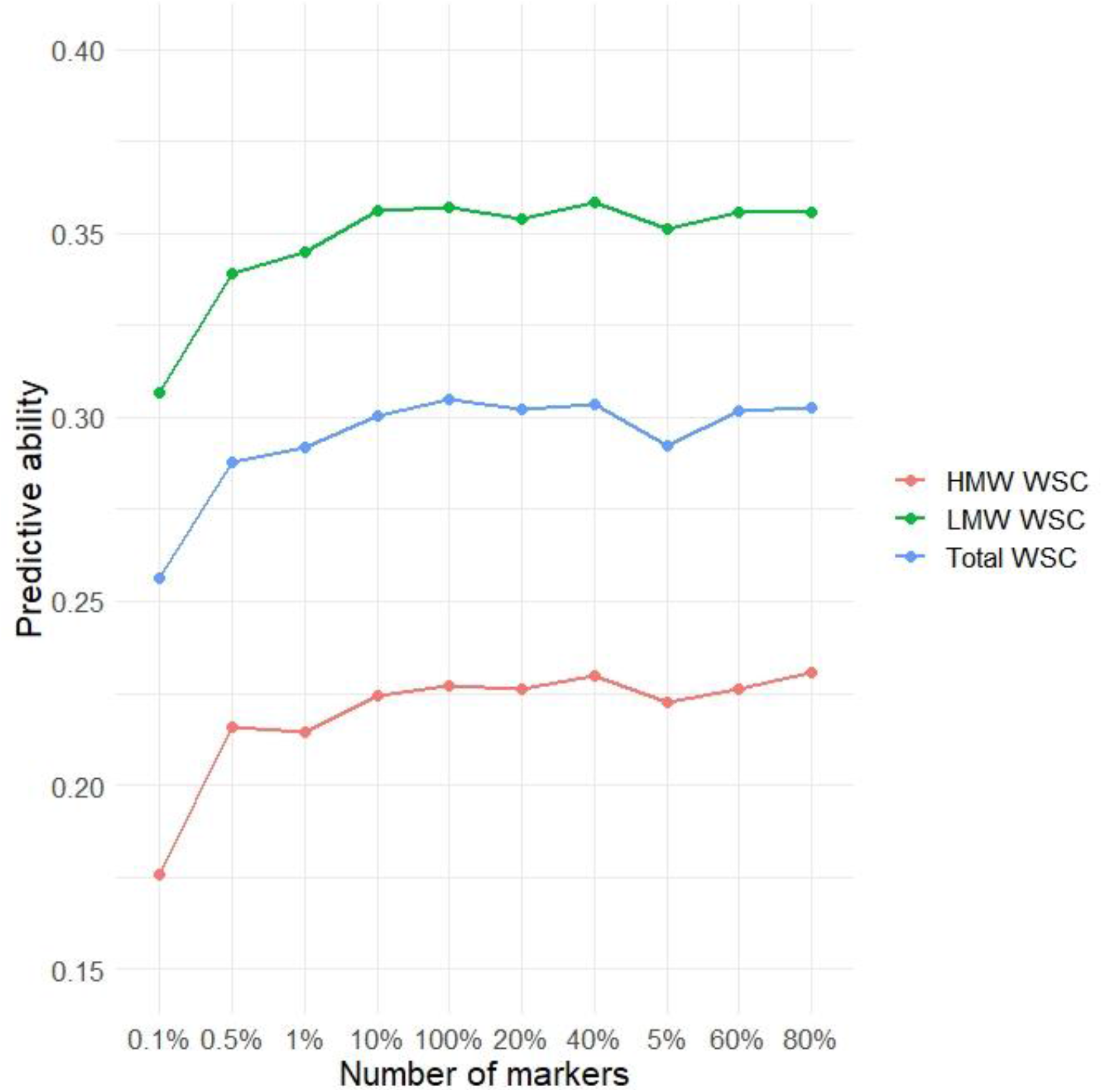
Random subsets of markers ranging from 0.1% (1,093) to 100% (1,093,464) of the marker set, used in GBLUP model to estimate predictive ability for HMW WSC, LMW WSC and Total WSC.

The training population used in this study is a composite of five different breeding populations, with differing genetic relationships (see Figure 1 in FAVILLE *et al.* (2018). The predictive ability of a model, constructed based on a composite training set, for each of the individual populations is therefore an important consideration. Cross-validations were conducted within the individual populations using the protocol reported by FAVILLE *et al.* (2018). Predictive ability varied amongst the populations (Figure 3). For example, predictive ability for ADF ranged from 0.13 to 0.24 amongst the five populations (Figure 3). The majority of predictions were positive across all populations, with the exception of K for Pop I, and only LMW WSC and P in Pop II had notably poor predictive ability (Figure 3). No population was superior for genomic prediction of all nutritive traits. However, Pop V returned the highest predictive ability overall (mean predictive ability of Pop V = 0.29, compared with 0.30 in the training set, TP), followed by Pop III, Pop I, Pop IV, and Pop II (Figure 3).

**Figure 3:**
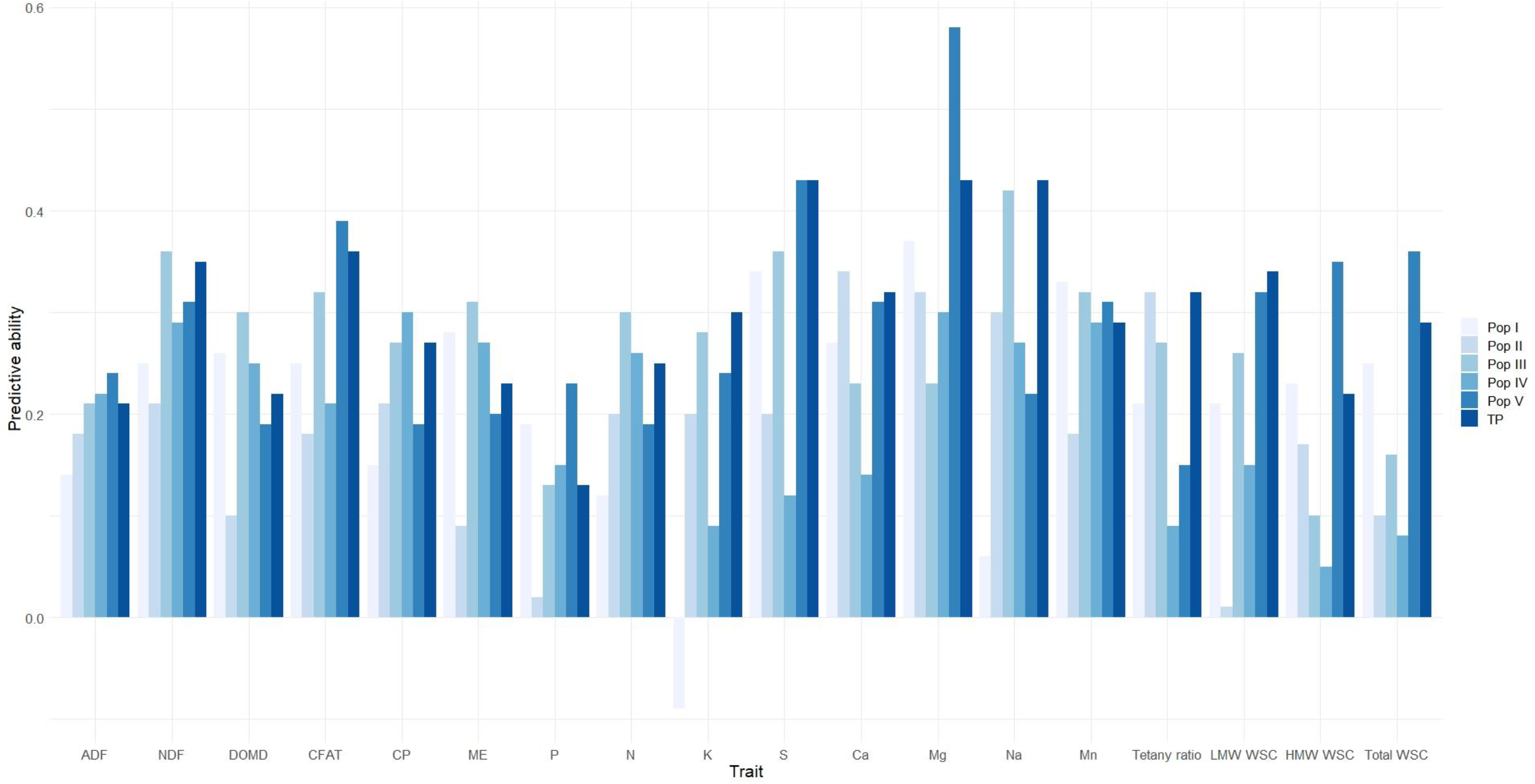
Predictive ability for 18 nutritive traits in each individual population (Pop I – Pop V) and in complete training population (TP)

## 4 Discussion

Nutritive quality traits in forages are important for animal productivity and for maintaining livestock health and are therefore important targets for genetic improvement in perennial ryegrass. Nutritive traits can be expensive to measure and are labour-intensive, hindering the improvement of these traits by conventional breeding methods. Genomic selection (GS), the use of genome-wide molecular markers for the prediction of breeding values in selection candidates, is well suited for traits that are costly and difficult to phenotype (HEFFNER *et al.* 2009; JANNINK *et al.* 2010) and therefore represents a promising approach for enabling cost-effective improvement of nutritive traits in forages. In this study we demonstrate that GS is a strong prospect for improvement of nutritive quality traits as assessed by cross-validation predictive abilities estimated for 18 nutritive traits in a multi-population training set. Furthermore, the extensive phenotypic dataset, collected from two contrasting environments, has enabled the contribution of genotypic, environment, and genotype-by-environment variance components to be estimated across a large range of nutritive traits.

Several methods for GS have been proposed for both plant and animal breeding, including GBLUP, Bayesian alphabets (BayesA, BayesB and BayesC), Ridge Regression (RR) BLUP, Random Forest, Support Vector Machine and deep learning through Multilayer Perceptron’s and Convolutional Neural Networks (DE LOS CAMPOS *et al.* 2013; CROSSA *et al.* 2017). Both simulations and empirical data suggests that linear models are superior in terms of predicting GEBVs at higher accuracy (DAETWYLER *et al.* 2010; DE LOS CAMPOS *et al.* 2013; BYRNE *et al.* 2017; BELLOT *et al.* 2018; FAVILLE *et al.* 2018). In this study, we compared three linear models characterized by two different assumptions with respect to the distribution of variance for marker effects. In GBLUP and KGD-GBLUP all marker effects are shrunk equally, assuming the predicted trait is controlled by many markers with small effect (GODDARD *et al.* 2011), whereas BayesC assumes that the trait is a mixture of distributions with large and small effect markers (HABIER *et al.* 2011). Even with different prior assumptions, Figure 1 illustrates the similarity in predictive ability amongst the three methods for all nutritive traits, with only minor differences (Figure 1). Through simulation and empirical data, DE LOS CAMPOS *et al.* (2013) pointed out that the superiority of Bayesian variable selection models can be illustrated when applied to a trait with large effect quantitative trait loci (QTL). The lack of improvement in predictive ability under the BayesC model observed here may reflect a complex genetic architecture for the nutritive traits studied, which are likely controlled by many genes with small effects. For instance, QTL studies in perennial ryegrass reported 25 loci for WSC (COGAN *et al.* 2005; TURNER *et al.* 2006; SHINOZUKA *et al.* 2012; GALLAGHER *et al.* 2015), however genetic variation explained by the multiple QTLs was no more than 20%, suggesting that genetic control of WSC may tend towards an infinitesimal model.

The success of GS primarily depends on the predictive ability of the genomic prediction model, which is influenced by 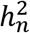, training population size, linkage disequilibrium (LD), genetic diversity within the training population and relatedness between training and test set (DAETWYLER *et al.* 2013; CROSSA *et al.* 2017; AROJJU *et al.* 2018). Traits with low 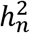 need a larger training population to achieve the same level of predictive ability as a trait with higher 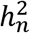. Results from our study indicate that predictive ability estimated by cross-validation and 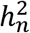 will not be a limiting factor for implementing GS for nutritive traits in perennial ryegrass, as predictive ability and various measures of heritability (*R*, 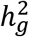 and 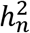) were moderate to high for most traits (Table 1, Supplementary Table S1-S7 and Figure 1). A strong positive correlation was observed between predictive ability and *h*^*2*^_*g*_ for traits at the individual locations (Aorangi and Lincoln) and in the across-location analysis, confirming previous findings (CROSSA *et al.* 2017) and suggesting that genomic prediction can be more accurate for highly heritable traits. A positive correlation between predictive ability and heritability was also previously observed for nutritive traits in switchgrass (FIEDLER *et al.* 2018) and alfalfa (JIA *et al.* 2018), as well as for crown rust and heading date in perennial ryegrass (AROJJU *et al.* 2018) and for fruit quality traits in apple (MURANTY *et al.* 2015).

For most traits, 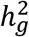 at Aorangi was consistently higher compared to Lincoln, and consequently higher predictive abilities were observed. This difference between locations was due to a combination of the genotypic variance component estimated at Aorangi being higher and estimates of trait-associated experimental error being higher at Lincoln (Supplementary Table S1 and S2). While it is not possible to conclusively determine the basis of this disparity in experimental error, it may be explained by greater within-environment variability at Lincoln, due to factors that such as climatic variations over the sampling period (Figure S1 in FAVILLE *et al.* (2018)), soil heterogeneity or operator-to-operator variations.

In contrast to switchgrass (FIEDLER *et al.* 2018) and alfalfa (JIA *et al.* 2018), prior to this study, genomic predictive ability for nutritive traits has been evaluated in perennial ryegrass for limited set of traits. FÈ *et al.* (2016) reported high predictive abilities of 0.68 for NDF and 0.45 for fructan in a large training set of 1918 F2 families, evaluated at multiple environments. In another study, GRINBERG *et al.* (2016) reported similarly high predictive abilities for WSC (0.59), DMD (0.41) and N (0.31) from prediction models applied in F14 generation families after training using a set of 364 families from earlier generations, phenotyped at a single location. Predictive ability for nutritive traits in the present study were overall lower compared to those reported by FÈ *et al.* (2016) and GRINBERG *et al.* (2016) with predictive abilities of 0.35, 0.29 and 0.22 for NDF, total WSC and HMW WSC (fructan), respectively. The lower predictive ability was likely affected by the smaller training population used in this study compared to FÈ *et al.* (2016), as well as its composite nature. Although, GRINBERG *et al.* (2016) reported high predictive ability for nutritive trait models, these values were based on a single environment and therefore unaffected by G × E, which might decrease the reliability of predictions. Overall, the values in the current study, based on a relatively small, composite training set were sufficiently high to support prediction of GEBVs and implementation of genomic selection to accelerate genetic gain for nutritive traits across environments in perennial ryegrass.

Determining the magnitude and genetic basis of G × E interactions for a trait is important, as it can assist in making appropriate breeding design decisions for the development of cultivars that are adapted to a broad range of target environments. In the current study G × E interactions were significant for all nutritive quality traits. The majority of traits displayed a G × E variance component that was small in comparison to genotypic variance, when nutritive traits were evaluated at two distinct locations (Table 1). This was reflected in the ratio of σ_g_ to σ_gs_, which was > 1 for 60% of the traits, indicating that the genotypic variance was predominant. However, the ratio for CFAT, CP, total WSC, LMW WSC, HMW WSC, P and N were < 1, indicating a greater influence of G × E interactions. The identification of high G × E interactions for WSC contrasts with results reported by EASTON *et al.* (2009) and are at variance with propositions by CASLER AND VOGEL (1999) and JAFARI (2012), that G × E for WSC are minimal to negligible. Our results are based on relatively large populations of half-sib families, compared to previous studies and may therefore be a more accurate reflection of the influence of G × E on these traits, particularly in New Zealand environments. However, it should be noted that the G × E interactions estimated here were based on only two locations, and a more robust estimation would be derived if based on a larger number of locations, representing the full target population of environments. The presence of G × E interactions may negatively influence ability to improve these traits for broad adaptation and represents a challenge during selections (HOLLAND *et al.* 2003).

Where G × E effects are large and significant, genetic improvement for a trait may only be achieved through selection based on multi-year, multi-environment evaluation. Considering the relatively high costs associated with phenotyping of nutritive quality traits, this approach might not always be feasible, and decisions will be based on available resources. Genomic selection, however, represents a promising approach to more directly tackle G × E. Models such as marker-by-environment interactions proposed by LOPEZ-CRUZ *et al.* (2015) and further developed by CROSSA *et al.* (2016), can be used to identify genomic regions that are stable across environments and other regions that are associated with specific environments that contribute to G × E interactions. These marker effects can be fixed in GS models to assist the selection of stable genotypes. However, these models were primarily developed for wheat, and a detailed investigation is needed to assess models perform in outcrossing species such as perennial ryegrass.

Traits with high G × E interactions displayed both lower 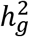 and comparatively low predictive abilities (Table 1, Supplementary Table S1-S2 and Figure 1). For such traits multi-trait genomic prediction models (JIA AND JANNINK 2012) may be one way of improving predictive ability and thereby genetic gain. The concept of multi-trait genomic prediction approaches is to improve the predictive ability of a primary target trait (which may be difficult and expensive to phenotype) by utilizing the genetic correlation with a secondary trait which is highly heritable and significantly less expensive to phenotype. Heritability and genotypic correlation data generated in the current study may assist in designing multi-trait prediction models for key nutritive traits. For example, a negative genetic correlation was observed between fibre and WSC traits, as reported previously in Italian ryegrass (WANG *et al.* 2015), and a positive genetic correlation was observed between DOMD and WSC traits as described previously by HUMPHREYS (1989b); JAFARI *et al.* (2003b) (Table 4). These secondary traits (ADF, NDF and DOMD) are measured routinely and relatively inexpensively by NIRS and may therefore be useful in multi-trait genomic prediction models to more accurately predict WSC traits that are most accurately measured using more expensive wet chemistry methodologies.

Mineral composition of forages is of interest from a perspective of livestock health and, as with nutritive traits overall, there has been little or no emphasis on selection for mineral composition in forage breeding programs (MASTERS *et al.* 2019). Significant genotypic variation was observed for all minerals in this study, with relatively low influence of G × E, moderate to high heritability and genomic prediction models with predictive abilities high in comparison to the other nutritive quality traits assessed (Figure 1). This indicates that selective breeding for levels of micro- and macro-minerals is feasible and that genomic selection represents a strong option for pursuing improvement in these traits. In general, ryegrass cultivars that grow well under low soil P will compete less for P in the sward, increasing P availability for uptake to support legume growth (EASTON *et al.* 1997; MCDOWELL *et al.* 2011). For instance, CRUSH *et al.* (2006), reported that in a mixed sward of ryegrass and clover (18% clover content), net annual flux of P into ryegrass was 4.7 times higher compared to clover. A small improvement in ryegrass phosphate use efficiency (PUE), can significantly change these proportions and may have large environmental and economic benefits (CRUSH *et al.* 2018a). In the current dataset predictive ability for P was very low (0.13), underpinned by a significant G × E interaction component to total phenotypic variation. This indicates that breeding for this P levels in perennial ryegrass foliage needs to be designed to account for G × E interaction effects. Alternatively, moderate to high genetic correlation with high 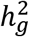 traits, such as Mg (genotypic correlation −0.62), might support an indirect multi-trait genomic selection strategy, as discussed earlier.

Hypomagnesaemia or grass tetany is a metabolic disorder in ruminants, caused by inadequate supply of Ca and Mg. This is often described in terms of a tetany index ([K/Ca+Mg]), for which values exceeding 2.2 (KEMP AND T HART 1957) are associated with increased risk of the disorder. We observed a moderate predictive ability for the ratio and the magnitude of G × E was low compared to genotypic variation, suggesting that tetany ratio could be used successfully as a selection criteria for developing cultivars with reduced potential for the incidence of hypomagnesaemia. This is in contrast to the results of SMITH *et al.* (1999), who reported large G × E variance for the tetany ratio evaluated at two locations in Australian environments and suggested the use Mg alone as a selection criteria to improve tetany ratio. Results from the current study showed a high predictive ability for Mg, making genomic selection a viable strategy for this trait. Although, increasing Mg concentration alone may be sufficient to decrease the incidence of hypomagnesaemia, the presence of a positive correlation between Mg and K observed in the current study (Table 2) and reported by SMITH *et al.* (1999), suggests that selections based on Mg concentration alone should be monitored and might not always give the expected outcome.

Using approximately 50k random markers the predictive ability of genomic prediction models for all nutritive traits was similar to using the full dataset of ca. 1M markers (Figure 2 and Supplementary Table S9). This reflects observations made in the same training set for herbage accumulation, a proxy for DM yield (Faville et al. 2018) except in that instance the marker subsets were not selected randomly. Below the 50k marker number there was a decrease in predictive ability, and this was particularly evident for traits with low 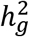. Considering the low levels of LD (*r*^*2*^ decaying to 0.25 after 366-1750 base pairs (FAVILLE *et al.* 2018)) observed in the component populations of the training set, the major proportion of predictive ability is likely a result of capturing relationship among individuals, rather than historical LD with QTL. In perennial ryegrass, to capture genetic variance associated with all causative QTL a very large number of markers and a large training population are needed, due to rapid decay of LD as a result of a very large past effective population size (*N*_*e*_) (HAYES *et al.* 2013; FIEDLER *et al.* 2018). Predictive ability based on relatedness between training and selection population can deteriorate after a few selection cycles (HABIER *et al.* 2007), and to maintain adequate predictive ability, either the training population should be very large and highly diverse or some form of relatedness should exit between training and selection population (HAYES *et al.* 2013; NORMAN *et al.* 2018).

In conclusion, genotypic variation and G × E interactions were significant for all nutritive quality traits evaluated in two distinct New Zealand environments. The predictive ability of genomic prediction models reported in this study for most of the traits would be sufficient to implement GS for nutritive traits in perennial ryegrass. Although a major proportion of this predictive ability is the result of capturing relatedness among individuals, maintaining relatedness between training and selection population would be an option to implement GS in perennial ryegrass. Predictive ability for most of the nutritive traits was retained even with as few as 50,000 markers. A next step would be to simulate a cost-benefit analysis to study the implications of manipulating marker number for cost-effective GS. For traits with low G × E interactions, single-trait genomic prediction models can be considered and for traits with large G × E, and consequently lower predictive ability, multi-trait approaches may be useful to explore as a method for obtaining high levels of prediction. This appears to be particularly important for WSC, which is considered to be one of the primary constituents of nutritive value for forages.

## 6 Author Contributions

MF, ZJ and BB conceived, designed, and coordinated the study. SKA, MC and MF performed the analysis. SKA and MF drafted the initial manuscript. SKA, MC, MJ, ZJ and BB contributed to interpretation of results and preparation of the final manuscript. All authors read and approved the final version.

## 8 Acknowledgments

The authors thank Dr. Alan Stewart, Keith Saulsbury and Tom Lyons of Grasslands Innovation Ltd for providing accesses to the ryegrass populations used in this study. We are grateful to Jason Trethewey, Anthony Hilditch, Lee Sutherland, Ivan Baird, Keith Widdup, Phil Rolston, Casey Flay, Jana Schmidt, Won Hong, Alieu Sartie, Aimee Avery, Anna Larking, Prue Taylor, Michelle Ebbett, Craig Anderson, Jessica O’Connor and many others for their assistance in trial harvests and sample processing. Special thanks to Dr. Andrew Griffith, Dr. Marcelo Carena and Dr. Valerio Hoyos-Villegas for their suggestions to improve this manuscript.

## 9 Supplementary Material

**Supplementary Table 1:** Trait genotypic (σ^2^_g_) and residual error (σ^2^_ε_) variance components, standard errors (SE), repeatability (*R*) and genomic heritability (*h*^*2*^_*g*_) and predictive ability (r_p_) estimated for 18 nutritive traits, among 517 half-sib families of perennial ryegrass evaluated at Lincoln.

**Supplementary Table 2:** Trait genotypic (σ^2^_g_) and residual error (σ^2^_ε_) variance components and their associated standard errors (SE), repeatability (*R*) and genomic heritability (*h*^*2*^_*g*_) and predictive ability (r_p_) estimated for 18 nutritive traits, among 517 half-sib families of perennial ryegrass evaluated at Aorangi.

**Supplementary Table 3:** Trait mean, standard deviation (σ), variance of genotype (σ^2^_g_), genotype-by-location interaction (σ^2^_gl_) and residual error (σ^2^_ε_), along with their associated standard errors (SE), and narrow-sense heritability (*h*^*2*^) estimated for a range of nutritive quality traits in Pop I (96 half-sib families), using data from across locations (Lincoln and Aorangi).

**Supplementary Table 4:** Trait mean, standard deviation (σ), variance of genotype (σ^2^_g_), genotype-by-location interaction (σ^2^_gl_) and residual error (σ^2^_ε_), along with their associated standard errors (SE) and narrow-sense heritability (*h*^*2*^) estimated for a range of nutritive quality traits in Pop II (110 half-sib families), using data from across locations (Lincoln and Aorangi).

**Supplementary Table 5:** Trait mean, standard deviation (σ), variance of genotype (σ^2^_g_), genotype-by-location interaction (σ^2^_gl_) and residual error (σ^2^_ε_), along with their associated standard errors (SE), and narrow-sense heritability (*h*^*2*^) estimated for a range of nutritive quality traits in Pop III (115 half-sib families), using data from across locations (Lincoln and Aorangi).

**Supplementary Table 6:** Trait mean, standard deviation (σ), variance of genotype (σ^2^_g_), genotype by location interaction (σ^2^_gl_) and residual error (σ^2^_ε_), along with their associated standard errors (SE) and narrow-sense heritability (*h*^*2*^) estimated for a range of nutritive quality traits in Pop IV (90 half-sib families), using the data from across locations (Lincoln and Aorangi).

**Supplementary Table 7:** Trait mean, standard deviation (σ), variance of genotype (σ^2^_g_), genotype by location interaction (σ^2^_gl_) and residual error (σ^2^_ε_), along with their associated standard errors (SE) and narrow-sense heritability (*h*^*2*^) estimated for a range of nutritive quality traits in Pop V (106 half-sib families), using data from across locations (Lincoln and Aorangi).

**Supplementary Table 8:** Phenotypic correlation for a range of nutritive quality traits among 517 half-sib families, estimated using data from across two locations (Lincoln and Aorangi).

**Supplementary Table 9:** Random subsets of markers ranging from 0.10% (n = 1,093) to 100% (n = 1,093,464) of the full GBS SNP dataset used in a GBLUP model to estimate predictive ability (r_p_) and bias (β) for 18 nutritive traits.

## 10 Data Availability Statement

The raw datasets used for the analysis or generated in the current study to draw conclusions will be made available upon request to any qualified researcher.

