## Supplementary Tables for "Genomic predictive ability for foliar nutritive traits in perennial ryegrass"

### 1 Supplementary Tables

**Supplementary Table 1:** Trait genotypic ($\sigma^2_g$) and residual error ($\sigma^2_\varepsilon$) variance components, standard errors (SE), repeatability (*R*) and genomic heritability ($h^2_g$) and predictive ability ($r_p$) estimated for 18 nutritive traits, among 517 half-sib families of perennial ryegrass evaluated at Lincoln.

| Trait | Abbreviation | $\sigma^2_g \pm$ SE | $\sigma^2_\varepsilon \pm$ SE | *R* | $h^2_g$ | $r_p$ |
|---|---|---|---|---|---|---|
| Acid detergent fibre | ADF | 0.49 ± 0.126 | 3.27 ± 0.162 | 0.31 | 0.31 | -0.06 |
| Neutral detergent fibre | NDF | 0.57 ± 0.142 | 3.64 ± 0.179 | 0.31 | 0.39 | 0.02 |
| Digestible organic matter in dry-matter | DOMD | 0.55 ± 0.168 | 4.63 ± 0.229 | 0.25 | 0.22 | -0.02 |
| Crude fat | CFAT | 0.02 ± 0.003 | 0.05 ± 0.003 | 0.51 | 0.56 | 0.25 |
| Metabolisable energy | ME | 0.01 ± 0.004 | 0.12 ± 0.005 | 0.25 | 0.23 | -0.02 |
| Crude protein | CP | 0.23 ± 0.098 | 2.94 ± 0.144 | 0.19 | 0.33 | 0.14 |
| Calcium | Ca | 0.96 ± 0.119† | 1.85 ± 0.094† | 0.6 | 0.69 | 0.34 |
| Potassium | K | 0.01 ± 0.003 | 0.07 ± 0.003 | 0.37 | 0.45 | 0.17 |
| Magnesium | Mg | 0.11 ± 0.015† | 0.27 ± 0.014† | 0.53 | 0.65 | 0.33 |
| Manganese (mg/kg) | Mn | 100.3 ± 20 | 454.3 ± 22.8 | 0.39 | 0.49 | 0.22 |
| Sodium | Na | 3.92 ± 0.460† | 6.64 ± 0.340† | 0.63 | 0.7 | 0.35 |
| Phosphorus | P | 0.11 ±0.039† | 1.11 ± 0.055† | 0.23 | 0.31 | 0.02 |
| Sulphur | S | 0.41 ± 0.074† | 1.58 ± 0.080† | 0.43 | 0.53 | 0.20 |
| Nitrogen | N | 5.56 ± 2.40† | 72.3 ± 3.50† | 0.18 | 0.31 | 0.14 |
| Tetany ratio (K/Ca+Mg) | Tetany ratio | 0.02 ± 0.002 | 0.03 ± 0.001 | 0.59 | 0.61 | 0.29 |
| Total water soluble carbohydrates | Total WSC | 142.8 ± 33.1 | 806.8 ± 40.3 | 0.33 | 0.39 | 0.08 |
| Low molecular weight carbohydrates | LMW | 34.2 ± 13.3 | 406 ± 19.6 | 0.2 | 0.12 | -0.03 |
| High molecular weight carbohydrates | HMW | 52.2 ± 13.1 | 329.3 ± 16.4 | 0.31 | 0.43 | 0.13 |

† $\times 10^{-3}$

**Supplementary Table 2:** Trait genotypic ($\sigma^2_g$) and residual error ($\sigma^2_\varepsilon$) variance components and their associated standard errors (SE), repeatability (*R*) and genomic heritability ($h^2_g$) and predictive ability ($r_p$) estimated for 18 nutritive traits, among 517 half-sib families of perennial ryegrass evaluated at Aorangi.

| **Trait** | **$\sigma^2_g$ ± SE** | **$\sigma^2_\varepsilon$ ± SE** | ***R*** | **$h^2_g$** | **$r_p$** |
|---|---|---|---|---|---|
| ADF | 0.60 ± 0.075 | 1.29 ± 0.061 | 0.56 | 0.65 | 0.29 |
| NDF | 0.86 ± 0.112 | 2.06 ± 0.097 | 0.54 | 0.64 | 0.35 |
| DOMD | 0.83 ± 0.130 | 2.80 ± 0.131 | 0.46 | 0.53 | 0.25 |
| CFAT | 0.01 ± 0.001 | 0.02 ± 0.001 | 0.59 | 0.69 | 0.33 |
| ME | 0.02 ± 0.003 | 0.07 ± 0.003 | 0.46 | 0.54 | 0.25 |
| CP | 0.37 ± 0.064 | 1.48 ± 0.069 | 0.42 | 0.59 | 0.21 |
| Ca | 0.60 ± 0.082† | 1.58 ± 0.074† | 0.52 | 0.6 | 0.16 |
| K | 0.01 ± 0.002 | 0.04 ± 0.002 | 0.46 | 0.6 | 0.28 |
| Mg | 0.18 ± 0.021† | 0.36 ± 0.017† | 0.59 | 0.69 | 0.34 |
| Mn | 53.6 ± 9.1 | 206.8 ± 9.7 | 0.41 | 0.57 | 0.24 |
| Na | 1.28 ± 0.018† | 3.44 ± 0.160† | 0.5 | 0.62 | 0.26 |
| P | 0.09 ± 0.014† | 0.03 ± 0.014† | 0.49 | 0.58 | 0.24 |
| S | 0.05 ± 0.064† | 1.03 ± 0.049† | 0.61 | 0.67 | 0.47 |
| N | 0.01 ± 0.001 | 0.03 ± 0.002 | 0.43 | 0.6 | 0.20 |
| Tetany ratio | 0.02 ± 0.002 | 0.04 ± 0.002 | 0.5 | 0.59 | 0.18 |
| Total WSC | 103.7 ± 12.9 | 226.5 ± 10.7 | 0.57 | 0.65 | 0.46 |
| LMW WSC | 49 ± 7.5 | 157.8 ± 7.3 | 0.48 | 0.59 | 0.33 |
| HMW WSC | 21.71 ± 2.3 | 33.11 ± 1.57 | 0.66 | 0.68 | 0.56 |

† $x10^{-3}$

**Supplementary Table 3:** Trait mean, standard deviation (σ), variance of genotype ($\sigma^2_g$), genotype-by-location interaction ($\sigma^2_{gl}$) and residual error ($\sigma^2_\varepsilon$), along with their associated standard errors (SE), and narrow-sense heritability (*h*²) estimated for a range of nutritive quality traits in Pop I (96 half-sib families), using data from across locations (Lincoln and Aorangi).

| **Trait** | $\sigma^2_g$ **± SE** | $\sigma^2_{gl}$ **± SE** | $\sigma^2_\varepsilon$ **± SE** | ***h²*** |
|---|---|---|---|---|
| ADF | 0.06±0.122 ‡ | 0.32±0.141 | 1.10±0.101 | 0.15 |
| NDF | 0.26±0.144 ‡ | 0.11±0.141 ‡ | 1.62±0.157 | 0.44 |
| DOMD | 0.17±0.156 ‡ | - | 2.72±0.228 | 0.27 |
| CFAT | 0.47±0.30† ‡ | 0.51±0.34 †‡ | 3.09±0.290† | 0.38 |
| ME | 0.53±0.4† ‡ | - | 7.01±0.590 | 0.31 |
| CP | 0.08±0.118 ‡ | 0.09±0.093 ‡ | 1.65±0.153 | 0.19 |
| Ca | 0.07±0.02† | 0.01±0.01 †‡ | 0.16±0.010† | 0.69 |
| K | 0.48±0.37† ‡ | 0.35±0.32 †‡ | 5.67±0.490† | 0.30 |
| Mg | - | - | - | 0.67 |
| Mn | 78.28±22.55 | 23.02±13.17 ‡ | 174.30±16.002 | 0.66 |
| Na | 0.11±0.03† | - | 0.33±0.030† | 0.67 |
| P | - | - | - | 0.55 |
| S | 0.04±0.01† | - | 0.09±0.010† | 0.73 |
| N | 0.02±0.22† ‡ | 0.39±0.28 †‡ | 4.2±0.390† | 0.02 |
| Tetany ratio | 0.10±0.0276 | - | 0.27±0.023 | 0.68 |
| Total WSC | 12.66±37.23 ‡ | 63.53±39.08 ‡ | 439.98±42.30 | 0.11 |
| LMW WSC | 11.66±7.641 ‡ | 14.21±6.77 | 84.71±7.705 | 0.35 |
| HMW WSC | 12.49±18.83‡ | - | 263.20±23.51 | 0.22 |

† $x10^{-2}$

‡ non-significant at the 0.05 probability level

**Supplementary Table 4:** Trait mean, standard deviation (σ), variance of genotype ($\sigma^2_g$), genotype-by-location interaction ($\sigma^2_{gl}$) and residual error ($\sigma^2_\varepsilon$), along with their associated standard errors (SE) and narrow-sense heritability ($h^2$) estimated for a range of nutritive quality traits in Pop II (110 half-sib families), using data from across locations (Lincoln and Aorangi).

| Trait | $\sigma^2_g$ ± SE | $\sigma^2_{gl}$ ± SE | $\sigma^2_\varepsilon$ ± SE | $h^2$ |
|---|---|---|---|---|
| ADF | 0.18±0.105 [‡] | 0.27±0.106 | 1.36±0.101 | 0.34 |
| NDF | - | - | - | - |
| DOMD | 0.24±0.190[‡] | 0.23±0.181[‡] | 3.74±0.274 | 0.25 |
| CFAT | 0.39±0.16[†] | 0.07±0.15[†‡] | 2.0±0.20[†] | 0.45 |
| ME | 0.61±0.49 [†‡] | 0.62±0.480[† ‡] | 9.6±0.710[†] | 0.24 |
| CP | 0.22±0.105 | 1.06±0.103[‡] | 1.72±0.129 | 0.39 |
| Ca | 0.06±0.01[†] | 0.01±0.01 [†‡] | 0.17±0.01[†] | 0.66 |
| K | 1.00±0.37[†] | 0.44±0.35[† ‡] | 4.7±0.37[†] | 0.50 |
| Mg | - | - | - | 0.68 |
| Mn | 55.77±26.139[†] | 36.76±25.98[†‡] | 356.94±27.602[†] | 0.42 |
| Na | 0.28±0.05[†] | 0.04±0.03[† ‡] | 0.35±0.03[†] | 0.78 |
| P | - | - | - | - |
| S | 0.01±0.01[†] | 0.02±0.01[†‡] | 0.13±0.01[†] | 0.16 |
| N | 0.51±0.24[†] | 0.20±0.24[†‡] | 4.23±0.31[†] | 0.39 |
| Tetany ratio | 0.10±0.02 | 0.021±0.01[‡] | 0.24±0.01 | 0.68 |
| Total WSC | 38.58±17.13 | 49.5±15.8 | 188.85±14.1 | 0.41 |
| LMW WSC | 11.52±5.6 | 18.60±6.01 | 60.02±4.6 | 0.37 |
| HMW WSC | 10.2±6.18 [‡] | 14.66±5.42 | 94.94±6.9 | 0.31 |

[†] x$10^{-2}$

[‡] non-significant at the 0.05 probability level

**Supplementary Table 5:** Trait mean, standard deviation ($\sigma$), variance of genotype ($\sigma^2_g$), genotype-by-location interaction ($\sigma^2_{gl}$) and residual error ($\sigma^2_\varepsilon$), along with their associated standard errors (SE), and narrow-sense heritability ($h^2$) estimated for a range of nutritive quality traits in Pop III (115 half-sib families), using data from across locations (Lincoln and Aorangi).

| Trait | $\sigma^2_g$ ± SE | $\sigma^2_{gl}$ ± SE | $\sigma^2_\varepsilon$ ± SE | $h^2$ |
|---|---|---|---|---|
| ADF | 0.31±0.114 | 0.20±0.115[‡] | 1.55±0.112 | 0.46 |
| NDF | 0.6±0.145 | 0.21±0.102 | 1.25±0.090 | 0.68 |
| DOMD | 0.47±0.172 | 0.48±0.168 | 1.83±0.134 | 0.46 |
| CFAT | 0.31±0.37[†‡] | 1.9±0.46[†] | 3.57±0.26[†] | 0.17 |
| ME | 1.20±0.45[†] | 1.19±0.43[†] | 4.86±0.35[†] | 0.48 |
| CP | 0.13±0.109 [‡] | 0.26±0.133[‡] | 1.67±0.119[‡] | 0.25 |
| Ca | 0.05±0.01[†] | 0.04±0.01[†] | 0.12±0.01[†] | 0.56 |
| K | 0.94±0.36[†] | 1.13±0.37[†] | 3.97±0.28[†] | 0.43 |
| Mg | - | - | - | 0.45 |
| Mn | 63.15±17.524 | 17.78±14.215[‡] | 225.42±16.181 | 0.58 |
| Na | 0.24±0.04[†] | - | 0.40±0.03[†] | 0.78 |
| P | - | - | - | 0.17 |
| S | 0.05±0.01[†] | 0.03±0.01[†] | 0.09±0.01[†] | 0.62 |
| N | 0.30±0.27 [†‡] | 0.70±0.34[†] | 0.04±0.3[†] | 0.20 |
| Tetany ratio | 0.07±0.021 | 0.0581±0.019 | 2.10±1.59 | 0.55 |
| Total WSC | 43.3±23.526 [‡] | 66.62±26.461 | 306.46±22.197 | 0.34 |
| LMW WSC | 14.7±8.168 [‡] | 28.01±9.337 | 93.07±6.838 | 0.33 |
| HMW WSC | 13.39±8.539[‡] | 16.74±10.170[‡] | 135.39±9.845 | 0.30 |

[†] $\times 10^{-2}$

[‡] non-significant at the 0.05 probability level

**Supplementary Table 6:** Trait mean, standard deviation (σ), variance of genotype ($\sigma^2_g$), genotype by location interaction ($\sigma^2_{gl}$) and residual error ($\sigma^2_\varepsilon$), along with their associated standard errors (SE) and narrow-sense heritability ($h^2$) estimated for a range of nutritive quality traits in Pop IV (90 half-sib families), using the data from across locations (Lincoln and Aorangi).

| Trait | $\sigma^2_g$ ± SE | $\sigma^2_{gl}$ ± SE | $\sigma^2_\varepsilon$ ± SE | $h^2$ |
|---|---|---|---|---|
| ADF | 0.70±0.216 | - | 0.99±0.142 | 0.81 |
| NDF | 0.69±0.225 | - | 1.24±0.167 | 0.77 |
| DOMD | 1.59±0.683 | - | 1.53±0.244 | 0.86 |
| CFAT | - | - | - | - |
| ME | 3.60±1.62† | 0.72±0.82† | 3.68±0.62†‡ | 0.79 |
| CP | 0.34±0.229‡ | 0.056±0.151 | 1.24±0.227 ‡ | 0.60 |
| Ca | 0.03±0.02† ‡ | 0.03±0.02† | 0.1±0.01† ‡ | 0.49 |
| K | 1.31±0.65† | 0.78±0.55† | 2.22±0.38†‡ | 0.63 |
| Mg | - | - | - | 0.66 |
| Mn | 39.9±45.340†‡ | 93.06±46.412† | 175.98±28.674 †‡ | 0.35 |
| Na | 0.07±0.03† | 0.06±0.0† | 0.18±0.03†‡ | 0.52 |
| P | 0.01±0.01† | 0.01±0.01† | 0.03±0.01†‡ | 0.36 |
| S | 0.04±0.02† | 0.04±0.02† | 0.05±0.01† ‡ | 0.62 |
| N | 0.81±0.52†‡ | 0.10±0.31† | 2.95±0.51†‡ | 0.60 |
| Tetany ratio | 0.10±0.034 | - | 0.17±0.023 ‡ | 0.79 |
| Total WSC | 50.3±26.384 ‡ | 13.16±21.149 | 208.64±27.998‡ | 0.55 |
| LMW WSC | 18.62±16.381 ‡ | 19.63±14.479 | 135.25±18.456‡ | 0.37 |
| HMW WSC | 7.09±9.480‡ | 41.74±13.329 | 24.35±5.692 | 0.22 |

† $\times 10^{-2}$

‡ non-significant at the 0.05 probability level

**Supplementary Table 7:** Trait mean, standard deviation (σ), variance of genotype ($\sigma^2_g$), genotype by location interaction ($\sigma^2_{gl}$) and residual error ($\sigma^2_\varepsilon$), along with their associated standard errors (SE) and narrow-sense heritability ($h^2$) estimated for a range of nutritive quality traits in Pop V (106 half-sib families), using data from across locations (Lincoln and Aorangi).

| **Trait** | $\sigma^2_g$ **± SE** | $\sigma^2_{gl}$ **± SE** | $\sigma^2_\varepsilon$ **± SE** | $h^2$ |
|---|---|---|---|---|
| ADF | 0.21±0.089 | 0.17±0.072 | 1.03±0.081 | 0.46 |
| NDF | 0.52±0.125 | 0.12±0.065 ‡ | 1.14±0.089 | 0.67 |
| DOMD | 0.60±0.186 | 0.29±0.147 | 1.87±0.148 | 0.57 |
| CFAT | 0.89±0.33† | 1.03±0.32† | 2.81±0.23† | 0.48 |
| ME | 1.61±0.49† | 0.77±0.38† | 4.85±0.38† | 0.58 |
| CP | 0.03±0.099†‡ | 29.23±11.65† | 122.2±97.8† | 0.08 |
| Ca | 0.03±0.01† | 0.03±0.01† | 0.13±0.01† | 0.48 |
| K | 0.70±0.32† | 0.49±0.31† | 4.37±00.35†‡ | 0.42 |
| Mg | - | - | - | 0.72 |
| Mn | 57.35±17.813 | 31.39±15.714 | 171.39±13.872 ‡ | 0.56 |
| Na | 0.20±0.05† | 0.13±0.04† | 0.41±0.03† | 0.60 |
| P | - | - | - | 0.31 |
| S | 0.01±0.01† | 0.02±0.01† | 0.07±0.01†‡ | 0.36 |
| N | 0.09±0.23†‡ | 0.66±0.28† | 3.15±0.25† | 0.10 |
| Tetany ratio | 0.023±0.01 | 0.01±0.008 | 0.16±0.012‡ | 0.40 |
| Total WSC | 57.8±27.232 | 85.90±27.256 | 303.92±24.30 | 0.38 |
| LMW WSC | 12.5±9.713‡ | 29.65±11.309 | 123.6±10.040 | 0.26 |
| HMW WSC | 19.79±8.325 | 25.99±7.742 | 90.03±7.068 | 0.41 |

† $\times 10^{-2}$

‡ non-significant at the 0.05 probability level

**Supplementary Table 8:** Phenotypic correlation for a range of nutritive quality traits among 517 half-sib families, estimated using data from across two locations (Lincoln and Aorangi).

| | ADF | NDF | DOMD | CFAT | ME | CP | N | Ca | K | Mg | Mn | Na | P | S | Tetany | LMW WSC | HMW WSC | Total WSC |
|---|---|---|---|---|---|---|---|---|---|---|---|---|---|---|---|---|---|---|
| ADF | 1 | 0.57 | −0.59 | −0.35 | −0.59 | −0.43 | −0.40 | 0.14 | −0.14 | 0.26 | 0.26 | 0.12 | −0.10 | 0.03† | −0.24 | −0.39 | −0.21 | −0.36 |
| NDF | | 1 | −0.73 | −0.05† | −0.73 | −0.25 | −0.23 | 0.09 | 0.03† | 0.38 | 0.21 | 0.14 | −0.07† | 0.06† | −0.14 | −0.62 | −0.44 | −0.63 |
| DOMD | | | 1 | 0.06† | 1 | 0.31 | 0.28 | −0.24 | 0.13 | −0.48 | −0.38 | −0.20 | 0.07† | −0.09† | 0.34 | 0.57 | 0.36 | 0.56 |
| CFAT | | | | 1 | 0.05† | 0.44 | 0.44 | −0.05† | 0.32 | −0.08† | −0.21 | 0† | 0.16 | 0.01 | 0.24 | 0.02† | −0.20 | −0.11† |
| ME | | | | | 1 | 0.3 | 0.28 | −0.23 | 0.13 | −0.49 | −0.37 | −0.21 | 0.07† | −0.09† | 0.34 | 0.56 | 0.37 | 0.55 |
| CP | | | | | | 1 | 0.99 | 0† | 0.42 | −0.01† | −0.13 | −0.14 | 0.28 | 0.25 | 0.25 | −0.05† | −0.45 | -0.29 |
| N | | | | | | | 1 | 0.01† | 0.43 | 0† | −0.12 | −0.14 | 0.27 | 0.25 | 0.25 | −0.06† | −0.45 | −0.30 |
| Ca | | | | | | | | 1 | −0.16 | 0.36 | 0.4 | 0.19 | 0.16 | 0.2 | −0.77 | −0.10 | −0.19 | -0.17 |
| K | | | | | | | | | 1 | 0.02† | −0.02† | −0.41 | 0.2 | 0.33 | 0.67 | −0.14 | −0.31 | −0.27 |
| Mg | | | | | | | | | | 1 | 0.41 | 0.15 | 0.1 | 0.33 | −0.48 | −0.39 | −0.38 | -0.45 |
| Mn | | | | | | | | | | | 1 | −0.10 | 0.13 | 0.25 | −0.35 | −0.23 | −0.17 | −0.24 |
| Na | | | | | | | | | | | | 1 | 0.03† | 0.01† | −0.40 | −0.07† | −0.10 | -0.1† |
| P | | | | | | | | | | | | | 1 | 0.39 | 0.01† | −0.01† | −0.21 | −0.13 |
| S | | | | | | | | | | | | | | 1 | −0.01† | −0.19† | −0.31† | -0.29 |
| Tetany | | | | | | | | | | | | | | | 1 | 0.08 | 0.03 | 0.06† |
| LMW WSC | | | | | | | | | | | | | | | | 1 | 0.43 | 0.85 |
| HMW WSC | | | | | | | | | | | | | | | | | 1 | 0.83 |
| Total WSC | | | | | | | | | | | | | | | | | | 1 |

† non-significant at the 0.05 probability level

**Supplementary Table 9:** Random subsets of markers ranging from 0.10% (n = 1,093) to 100% (n = 1,093,464) of the full GBS SNP dataset used in a GBLUP model to estimate predictive ability ($r_p$) and bias (β) for 18 nutritive traits.

| Trait | 100% | | 80% | | 60% | | 40% | | 20% | |
|---|---|---|---|---|---|---|---|---|---|---|
| | $r_p$ | β | $r_p$ | β | $r_p$ | β | $r_p$ | β | $r_p$ | β |
| ADF | **0.22** | 0.98 | **0.22** | 1.02 | **0.22** | 0.99 | **0.22** | 1.00 | **0.22** | 1.00 |
| NDF | **0.37** | 0.94 | **0.36** | 0.90 | **0.37** | 0.92 | **0.37** | 0.94 | **0.36** | 0.92 |
| DOMD | **0.23** | 0.95 | **0.22** | 0.93 | **0.22** | 0.90 | **0.22** | 0.88 | **0.22** | 0.90 |
| CFAT | **0.35** | 1.02 | **0.36** | 1.04 | **0.35** | 1.01 | **0.36** | 1.03 | **0.35** | 1.02 |
| ME | **0.23** | 0.86 | **0.23** | 0.91 | **0.23** | 0.89 | **0.23** | 0.91 | **0.23** | 0.87 |
| CP | **0.27** | 1.00 | **0.26** | 0.98 | **0.27** | 1.00 | **0.26** | 0.98 | **0.27** | 0.99 |
| N | **0.25** | 1.01 | **0.25** | 1.01 | **0.25** | 0.99 | **0.25** | 1.02 | **0.25** | 1.01 |
| Ca | **0.31** | 0.98 | **0.31** | 0.97 | **0.31** | 0.95 | **0.31** | 0.99 | **0.30** | 0.97 |
| K | **0.32** | 1.01 | **0.32** | 1.02 | **0.32** | 1.00 | **0.32** | 1.02 | **0.32** | 1.00 |
| Mg | **0.45** | 1.02 | **0.45** | 1.03 | **0.45** | 1.02 | **0.44** | 1.01 | **0.44** | 1.02 |
| Mn | **0.31** | 1.03 | **0.31** | 1.01 | **0.31** | 1.01 | **0.31** | 1.01 | **0.30** | 1.00 |
| Na | **0.44** | 1.00 | **0.43** | 1.00 | **0.43** | 0.98 | **0.44** | 1.00 | **0.43** | 1.00 |
| P | **0.14** | 1.46 | **0.14** | 1.04 | **0.15** | 1.08 | **0.15** | 1.33 | **0.15** | 1.05 |
| S | **0.45** | 0.97 | **0.45** | 0.97 | **0.44** | 0.96 | **0.45** | 0.98 | **0.45** | 0.98 |
| Tetany ratio | **0.34** | 1.00 | **0.34** | 0.98 | **0.34** | 1.01 | **0.34** | 1.00 | **0.34** | 0.97 |
| LMW WSC | **0.36** | 1.04 | **0.36** | 1.04 | **0.36** | 1.04 | **0.36** | 1.04 | **0.35** | 1.04 |
| HMW WSC | **0.23** | 1.07 | **0.23** | 1.07 | **0.23** | 1.05 | **0.23** | 1.12 | **0.23** | 1.05 |
| Total WSC | **0.31** | 1.03 | **0.30** | 1.02 | **0.30** | 1.00 | **0.30** | 1.03 | **0.30** | 1.02 |

Continuation of table S9

| Trait | 10% | | 5% | | 1% | | 0.50% | | 0.10% | |
|---|---|---|---|---|---|---|---|---|---|---|
| | $r_p$ | β | $r_p$ | β | $r_p$ | β | $r_p$ | β | $r_p$ | β |
| ADF | **0.22** | 0.99 | **0.22** | 1.00 | **0.20** | 1.10 | **0.19** | 1.08 | **0.15** | 1.22 |
| NDF | **0.36** | 0.93 | **0.37** | 0.96 | **0.35** | 0.96 | **0.34** | 0.98 | **0.29** | 1.00 |
| DOMD | **0.22** | 0.92 | **0.22** | 0.92 | **0.21** | 0.95 | **0.20** | 0.98 | **0.15** | 1.40 |
| CFAT | **0.35** | 1.01 | **0.35** | 1.01 | **0.34** | 1.03 | **0.31** | 0.99 | **0.27** | 1.03 |
| ME | **0.23** | 0.92 | **0.23** | 0.90 | **0.22** | 1.05 | **0.20** | 0.95 | **0.16** | 1.11 |
| CP | **0.26** | 0.98 | **0.26** | 0.98 | **0.25** | 1.00 | **0.24** | 1.01 | **0.20** | 1.07 |
| N | **0.25** | 1.00 | **0.25** | 1.00 | **0.24** | 1.03 | **0.23** | 1.03 | **0.20** | 1.14 |
| Ca | **0.30** | 0.98 | **0.30** | 0.97 | **0.29** | 1.00 | **0.27** | 0.99 | **0.22** | 1.06 |
| K | **0.32** | 1.02 | **0.32** | 1.00 | **0.31** | 1.03 | **0.31** | 1.05 | **0.26** | 1.04 |
| Mg | **0.44** | 1.01 | **0.44** | 0.99 | **0.41** | 0.95 | **0.39** | 0.96 | **0.33** | 1.00 |
| Mn | **0.31** | 1.02 | **0.29** | 0.97 | **0.28** | 1.00 | **0.25** | 1.01 | **0.17** | 1.35 |
| Na | **0.44** | 1.00 | **0.43** | 0.98 | **0.41** | 0.98 | **0.40** | 0.99 | **0.34** | 1.01 |
| P | **0.14** | 0.99 | **0.14** | 1.20 | **0.14** | 1.04 | **0.13** | 1.67 | **0.09** | 1.46 |
| S | **0.45** | 0.96 | **0.45** | 0.98 | **0.43** | 0.96 | **0.42** | 0.98 | **0.37** | 0.99 |
| Tetany ratio | **0.34** | 0.98 | **0.33** | 0.99 | **0.32** | 1.00 | **0.31** | 1.00 | **0.26** | 1.02 |
| LMW WSC | **0.36** | 1.05 | **0.35** | 1.03 | **0.34** | 1.04 | **0.34** | 1.05 | **0.31** | 1.06 |
| HMW WSC | **0.22** | 1.05 | **0.22** | 1.04 | **0.21** | 1.04 | **0.22** | 1.13 | **0.18** | 1.26 |
| Total WSC | **0.30** | 1.02 | **0.29** | 0.99 | **0.29** | 1.02 | **0.29** | 1.04 | **0.26** | 1.04 |
